# Distrust Before First Sight? Examining Knowledge- and Appearance-Based Effects of Trustworthiness on the Visual Consciousness of Faces

**DOI:** 10.1101/2021.02.24.432562

**Authors:** Anna Eiserbeck, Alexander Enge, Milena Rabovsky, Rasha Abdel Rahman

**Author notes:** Correspondence concerning this preprint should be addressed to Anna Eiserbeck or Rasha Abdel Rahman, Humboldt-Universität zu Berlin, Institut für Psychologie, Unter den Linden 6, 10099 Berlin, Germany. or.

## Abstract

The present EEG study with 32 healthy participants investigated whether affective knowledge about a person influences the visual awareness of their face, additionally considering the impact of facial appearance. Faces differing in perceived trustworthiness based on appearance were associated with negative or neutral social information and shown as target stimuli in an attentional blink task. As expected, participants showed enhanced awareness of faces associated with negative compared to neutral social information. On the neurophysiological level, this effect was connected to differences in the time range of the early posterior negativity (EPN)—a component associated with enhanced attention and facilitated processing of emotional stimuli. The findings indicate that the social-affective relevance of a face based on emotional knowledge is accessed during a phase of attentional enhancement for conscious perception and can affect prioritization for awareness. In contrast, no clear evidence for influences of facial trustworthiness during the attentional blink was found.

## 1. Introduction

Of the wealth of sensory information available at any given moment in time, we consciously perceive only a fraction (see, e.g., Cohen et al., 2016; Sklar et al., 2021). More is yet to be learned about which factors exactly determine or modulate the extent to which we are consciously aware of the stimuli in our surroundings. In the present article, we examine the influences of social-emotional factors underlying trustworthiness attributions on the access of faces to visual awareness. Specifically, we focus on the effects of affective knowledge while also considering the impact of visual appearance.

### 1.1. Influences of Affective Knowledge on Evaluations and Face Processing

Previous research has demonstrated that in addition to the visual information itself, context plays a crucial role in face perception (for a review, see Wieser & Brosch, 2012). What we know about a person’s character and past actions can affect not only how we judge them and react to them in social situations but also how we see their face (for a review, see Maier et al., 2022). This influence of social-emotional information on face perception is reflected, for instance, in modulations of ratings of attractiveness, facial features, or emotional expressions (Hassin & Trope, 2000; Nisbett & Wilson, 1977; Paunonen, 2006; Suess et al., 2015).

On the neural level, differences based on affective information have been most consistently observed in two event-related potential (ERP) components, associated with early perceptual and later evaluative processing, respectively: the early posterior negativity (EPN; e.g., Baum & Abdel Rahman, 2021b, 2021a; Luo et al., 2016; Schindler et al., 2021; Suess et al., 2015; Wieser et al., 2014; Xu et al., 2016) and the late positive potential (LPP; e.g., Abdel Rahman, 2011; Baum et al., 2020; Baum & Abdel Rahman, 2021b, 2021a; Klein et al., 2015; Luo et al., 2016; Schindler et al., 2021; Xu et al., 2016). The EPN, a relative negativity occurring at around 200 to 300 ms after stimulus onset at occipito-temporal sites, has been linked to enhanced attention to and facilitated perceptual processing of emotional compared to more neutral stimuli (e.g., Junghöfer et al., 2001; Schacht & Sommer, 2009; Schupp et al., 2003, 2004). The LPP, a relative positivity between around 400 to 600 ms at centro-parietal sites, has likewise been found to be enhanced for emotional compared to neutral stimuli and is viewed as an index of more elaborate processing and higher-order evaluation of affective information (e.g., Schacht & Sommer, 2009; Schupp et al., 2004; for a review of EPN and LPP effects for emotional faces, see Schindler & Bublatzky, 2020). In addition to these two frequently observed effects, which are regarded as markers of the processing of emotional stimuli in general, research on the effects of affective information on face perception has also focused on the earlier N170 component (for a review, see Schindler et al., 2023). The N170, which is observable as a negative deflection between approximately 130 and 200 ms over occipitotemporal electrode sites, is regarded as a marker of the structural encoding and configural processing of facial information (Bentin et al., 1996; Eimer et al., 2010; Hinojosa et al., 2015). While evidence across different studies displays considerable variance, findings overall show increased N170 amplitudes for faces of emotional relevance, indicating an early prioritized processing of configural facial information (Schindler et al., 2023).

### 1.2. Influences of Facial Trustworthiness on Evaluations and Face Processing

A second factor determining face perception and person evaluation relates directly to facial appearance (for a review, see Todorov et al., 2015). Specifically, trustworthiness impressions based on facial features, so-called “facial trustworthiness,” represent a central dimension underlying evaluations that closely corresponds to the general perceived valence of faces with neutral expressions (Oosterhof & Todorov, 2008). Facial trustworthiness is typically measured by obtaining trustworthiness judgments from a group of participants for faces displaying a neutral expression and taking the mean as the perceived trustworthiness score of the respective face (see, e.g., Lischke et al., 2018; Shore et al., 2017; Verosky et al., 2018). Studies observed a high agreement in trustworthiness judgments among raters based on facial appearance (Oosterhof & Todorov, 2008; Rule et al., 2013). An underlying basis has been found to be emotion overgeneralization (Oosterhof & Todorov, 2009; Said et al., 2009; Zebrowitz & Montepare, 2008): Even when a person displays a neutral expression, facial features can lead to the impression of subtle emotional expressions. Specifically, facial trustworthiness judgments have been found to be sensitive to features resembling subtle approach or avoidance signals (Oosterhof & Todorov, 2008): Faces judged as trustworthy bear resemblances of happy expressions (e.g., mouth forms a gentle upward curve; eyes appear relaxed and wider open), while those judged as untrustworthy exhibit features reminiscent of anger (e.g., edges of the mouth curled down, eyebrows appear lower and drawn together).

Facial trustworthiness has been found to influence impressions and character judgments, even when explicit knowledge about the respective person is available (Todorov & Olson, 2008; Verosky et al., 2018), and facial trustworthiness and knowledge may interact with each other (Rule et al., 2012). Notably, it is highly questionable whether character inferences from appearance in general (Todorov et al., 2015) and trustworthiness evaluations in specific (see Introduction in Jaeger et al., 2022 for a review of evidence) possess any validity at all. A recent study (Jaeger et al., 2022) showed that participants’ trust decisions and trustworthiness predictions of strangers were unrelated to the actual trustworthiness of their counterparts. To a large degree such attributions are therefore likely to represent merely a type of appearance-based prejudice.

ERP effects of facial trustworthiness—like affective knowledge effects—are most consistently reported in the EPN time range (Dzhelyova et al., 2012; trustworthiness evaluation task in Marzi et al., 2014; Rudoy & Paller, 2009) and in the LPP time range (Lischke et al., 2018; trustworthiness evaluation task in Marzi et al., 2014; Rudoy & Paller, 2009; Yang et al., 2011). Yet, the overall evidence is quite heterogeneous, with some studies also finding differences in the earlier C1 (Yang et al., 2011), P1 (trustworthiness evaluation task in Marzi et al., 2014; neurotypical adults in Shore et al., 2017), or N170 components (Dzhelyova et al., 2012; political decision task in Marzi et al., 2014).

### 1.3. Social-Emotional Influences on Face Awareness: Current State of Evidence

The discussed evidence demonstrates robust knowledge-based and appearance-induced influences on person perception and evaluation. A currently unresolved question is whether conscious perception is necessary for integrating this information (for a review on processing without awareness, see Mudrik & Deouell, 2022) or whether the processing takes place already beforehand, having the potential to influence what we consciously perceive in the first place. Can social-affective knowledge, possibly in interaction with appearance, influence whether or to what extent we consciously perceive a face?

To investigate this question, two kinds of methods for suppressing stimulus awareness have been used: Perceptual suppression methods, namely binocular rivalry and continuous flash suppression, and an attentional suppression method, namely the attentional blink paradigm. In binocular rivalry studies (for a review, see Alais, 2012), a different stimulus is presented to each eye (dichoptic presentation), and conscious perception alternates between the images. Thus, binocular rivalry represents a suppression of low-level sensory signals due to interocular suppression. Continuous flash suppression (for a review, see Pournaghdali & Schwartz, 2020) represents a particular case of dichoptic presentation, where the presentation of fast-changing masks (so-called “Mondrians”) to one eye suppresses the perception of a target stimulus (e.g., an image of a face) presented to the other eye. In the attentional blink paradigm (Raymond et al., 1992), participants are instructed to detect two target stimuli—T1 and T2—among a series of distractor images in a Rapid Serial Visual Presentation (RSVP) stream. Successful detection of T1 thereby often impairs the detection of T2 when it follows in close temporal succession of approximately 200 to 500 ms (short lag), whereas detection is largely unimpaired for longer intervals (long lag). This attentional blink has been ascribed to disruptions during attentional engagement and/or memory encoding of T2 due to the ongoing processing of the T1 stimulus (for a review, see Zivony & Lamy, 2022). The attentional blink paradigm enables an investigation of the attributes of a stimulus that determine access to conscious perception when attentional resources are limited.

Concerning facial trustworthiness, some studies report effects on the access to visual consciousness, as reflected in a longer time to emerge from suppression in a continuous flash suppression task for untrustworthy as compared to neutral faces (Getov et al., 2015; Stewart et al., 2012). However, a subsequent study indicated that this effect may be explained by low-level visual differences (Stein et al., 2018). A further study (Abir et al., 2017) observed that a faster emergence of faces to awareness under continuous flash suppression is determined by a “priority dimension”, which correlated strongly and positively with perceived dominance scores and weaker and negatively with trustworthiness/valence scores (i.e., untrustworthy faces break faster into consciousness, contradicting the direction observed in the previous studies, Getov et al., 2015; Stewart et al., 2012). Yet, considering the correlation of the valence/trustworthiness scores with response times separately, in a control experiment a significant correlation was found in a conscious condition but not in the unconscious condition (see Figure 3 and experiment 6 in Abir et al., 2017), indicating that the effect of trustworthiness may be due to conscious rather than preconscious processing.

With respect to face-related affective knowledge, initial evidence for an effect on visual awareness came from a study using binocular rivalry (E. Anderson et al., 2011) in which faces previously associated with negative socially relevant information were found to dominate longer in visual consciousness than faces associated with positive or neutral information. However, this finding might not necessarily indicate prioritized access to consciousness since the measure of visual dominance could also reflect later conscious prioritization (see Stein et al., 2017), and indeed, no effect has been observed for the first percept to be reported. In subsequent studies using binocular rivalry or breaking continuous flash suppression, no evidence for an influence of affective knowledge on the access to visual consciousness was found (Rabovsky et al., 2016; Stein et al., 2017). Another study in which the affective value of faces was acquired through a different manipulation, namely monetary win and loss associations, also found no awareness effects of this factor in continuous flash suppression and attentional blink experiments (Stein & Verosky, 2021).

However, the findings of a recent study (Eiserbeck & Abdel Rahman, 2020) using an attentional blink paradigm (Raymond et al., 1992) did provide additional support for the hypothesis of an impact of social-affective knowledge on visual consciousness. In line with an often observed detection advantage for emotional stimuli in the attentional blink (e.g., A. K. Anderson & Phelps, 2001; De Martino et al., 2009; de Oca et al., 2012; Maratos et al., 2008; Milders et al., 2006; Schwabe et al., 2011), enhanced awareness was observed for faces associated with negative as compared to neutral social behavioral information, whereas no effect of facial trustworthiness was found (Eiserbeck & Abdel Rahman, 2020). Yet, the results of this study have left questions open: The null effect of facial trustworthiness might be due to the fact that this factor comprised only two levels— average-trustworthy and low-trustworthy faces. Effects might depend on the inclusion of a broader range from low-to high-trustworthy faces. Furthermore, no clear all-or-none pattern was found for influences of affective knowledge on visual consciousness, but rather a modulation of the strength or quality of the resulting percept—which raises the question of whether the differences occurred during a time of attentional enhancement for visual consciousness, or at a later point in time. Although the results are in line with accounts that assume graded consciousness in the attentional blink (e.g., Fazekas & Overgaard, 2018; also see Eiserbeck et al., 2022), more direct evidence on the time course of the processing of affective knowledge in regard to the access to conscious perception is needed. The high temporal resolution of the EEG and obtained event-related potentials may help to cast light on this matter.

### 1.4. Time Course of the Transition to Visual Awareness

One challenge with respect to investigating the neural processing of emotional influences on visual awareness consists in the fact that it has not yet been resolved exactly when and how the access to conscious awareness occurs. In ERP research, mainly two components have been considered as potential makers of visual consciousness (for reviews, see Förster et al., 2020; Koivisto & Revonsuo, 2010): The visual awareness negativity (VAN) and the late positivity (LP). The VAN is a relative negativity observable in the contrast between aware and unaware stimuli at posterior electrode sites. It can occur as early as 100 ms after stimulus onset and last up to about 350 ms, including the time span of the N1 and N2 components. The LP is a relative positivity typically observed around 300-600 ms after stimulus onset, in the time range of the P3 component, which has likewise often been observed in the contrast between aware and unaware stimuli. The VAN has been described as the correlate of visual awareness most consistently found across studies (Förster et al., 2020; Koivisto & Revonsuo, 2010), and it is in line with theories assuming that the subjective experience of seeing occurs already during early stages of visual processing, e.g., Recurrent Processing Theory (Lamme, 2010; Lamme & Roelfsema, 2000). Other perspectives (Global Neuronal Workspace Theory; Dehaene & Changeux, 2011; Dehaene & Naccache, 2001) suggest that modulations during the VAN time range reflect preconscious differences and that later activity occurring beyond 250 ms marks the access to consciousness. Specifically, the P3 component had been suggested as a potential marker of conscious access, based on evidence from visual masking (Del Cul et al., 2007) as well as the attentional blink (Sergent et al., 2005). However, recent evidence suggests that this component might be more strongly linked to processing following initial conscious perception (for a review, see Förster et al., 2020). Specifically, using no-report paradigms it was shown that the P3 response depended on the task relevance of the targets, as tested in experiments using visual masking (Cohen et al., 2020), a GO-NOGO-task (Koivisto et al., 2016), and inattentional blindness (Dellert et al., 2021; Pitts et al., 2014; Schlossmacher et al., 2020; Shafto & Pitts, 2015). A recent attentional blink study (Dellert et al., 2022), in which task relevance and awareness were manipulated independently, showed that conscious perception covaried with the VAN but not the P3. While these findings do not preclude that later activity might still be relevant in regard to the overall visual experience (e.g., enabling the extraction of more information due to enhanced processing in working memory), in sum, evidence points towards the VAN as the earliest, most consistent correlate of visual awareness.

Based on the above evidence, we assume that the subjective experience of a visual stimulus first arises during the time range of the VAN. Thereby, rather than an immediate conscious experience of all visual stimulus information, the conscious percept might evolve over time (Aru & Bachmann, 2017; Bachmann, 2019; Campana et al., 2016). For example, the very first time point during which VAN differences are observed may not indicate the experience of a complete and finalized percept. Rather, an initial coarse percept may be enriched by more detailed visual information over the course of processing as reflected by the VAN. This emergence of awareness may be modulated by specific factors, such as a stimulus’ emotional value (cf. Figure 1 in Aru & Bachmann, 2017). The early ERP correlates, i.e., the described EPN effects, of social-affective knowledge and visually derived trustworthiness fall within the time window of the VAN. In regard to the time course of processing, the possibility of an influence of these emotional aspects on visual awareness therefore appears plausible. Furthermore, the functional significance of the EPN component, taken to reflect enhanced attention towards and prioritized processing of certain stimuli, matches these considerations. Under conditions of overall reduced attention limiting what enters visual awareness, this emotionally-based enhanced processing could lead to enhanced visual awareness of respective stimuli. Thus, affective knowledge and facial trustworthiness may influence the visual awareness of faces.

**Figure 1.**
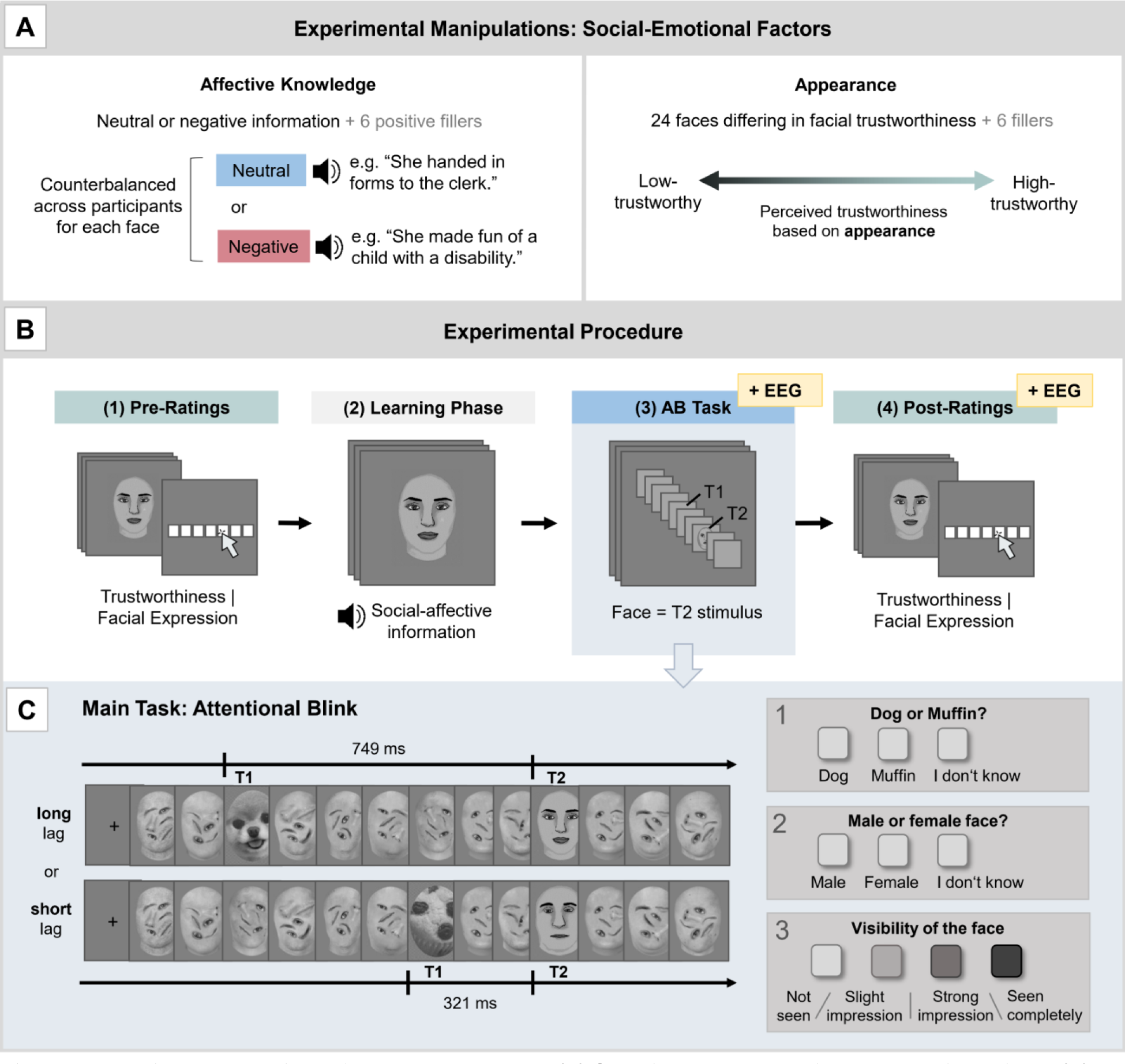
Experimental manipulations and procedure. (A) Overview of the experimental manipulations. (B) The procedure consisted of: (1) pre-ratings of trustworthiness and facial expression (with task order counterbalanced across participants), (2) learning of affective person knowledge, (3) the attentional blink task, and (4) post-ratings. (C) Illustration of T2-present trials in the attentional blink task. 13 images were shown in rapid succession, with a presentation time of 107 ms each. In long lag trials, there was a 749 ms interval between T1-onset and T2-onset. In short lag trials, there was a 321 ms interval. T2 faces differed in associated knowledge (neutral or negative, counterbalanced across participants) and facial trustworthiness. After each trial, participants answered three questions via button press regarding the identity of T1, the gender of the T2 face, and the visibility of T2. Please note: For illustration in this preprint, the face stimuli used in the experiment (edited photographs of real faces; see description in the Methods section) have been replaced by roughly similar looking drawings.

To summarize, the relationships between ERPs connected to visual awareness and emotional processing can be described as follows: The EPN and VAN both have been connected to enhanced (conscious) processing of stimuli, and they occur within similar time ranges and with similar topographies. The VAN component reflects overall differences between aware and unaware stimuli, whereas the EPN represents differences between emotional and neutral stimuli. Thus, while the VAN is associated with perceiving stimuli in general, the EPN is linked explicitly to enhanced processing of affective stimuli. The LP is a relative positivity during the P3 time range that is often observed in the contrast between detected and undetected stimuli. Yet, it may reflect post-perceptual processing influenced by task relevance rather than visual awareness per se. The LPP component is a relative positivity observable in the contrast between emotional and neutral stimuli, with a highly similar time range and topography as the LP. This emotion-specific activity reflected in the LPP has likewise been found to be modulated by task relevance (Schindler & Straube, 2020).

### 1.5. Present Study

Building on the described evidence, in the present event-related potential (ERP) study, we examined how the visual awareness of faces depends on social-emotional factors. We focused on the question of whether social-affective knowledge about a person affects visual awareness. In pursuing this question, we simultaneously controlled for and compared the influence of visually derived trustworthiness impressions based on facial appearance. As outlined above, facial trustworthiness represents perceptually salient affective information that may interact with social-affective knowledge. Taking into account and comparing the impact of facial trustworthiness as a second, more strongly visually based source of attributed affective value may be informative in regard to the mechanisms underlying access of faces to visual consciousness: Is visual consciousness influenced by the overall affective/trustworthiness value ascribed to a face or does it depend on the type of information? Which kind of information has a stronger impact, and do the different factors interact?

In the experiment, faces differing in facial trustworthiness (covering a range from low to high facial trustworthiness) were associated with negative or neutral social information, with manipulation checks for both factors included in the experiment. Subsequently, they were presented as T2-stimuli in an attentional blink task to investigate the effects of both factors on visual awareness. The EEG was tracked to examine the time course of the effects. As more closely described in the methods sections, awareness/unawareness of faces was determined through a combination of two criteria: subjective visibility ratings on a variant of a perceptual awareness scale (PAS; Ramsøy & Overgaard, 2004) and a more objective categorization task. In behavioral data, we expected higher awareness for faces associated with negative as compared to neutral knowledge (Eiserbeck & Abdel Rahman, 2020) and for less trustworthy as compared to more trustworthy-looking faces (Abir et al., 2017). Based on reported congruency effects of affective knowledge and facial trustworthiness (e.g., in memory: Rule et al., 2012), we furthermore expected an interaction between both factors, with enhanced awareness for faces with congruent negative information (negative knowledge combined with less trustworthy facial appearance).

ERP analyses for face processing in the attentional blink were based on an early time interval and posterior region of interest typical for both EPN and VAN and a later time interval and central region of interest typical for both LPP and LP. While the overlapping sets of components (EPN/VAN; LPP/LP) are not readily discernible based on their latency or topography, we distinguish them based on the aspects that they reflect, namely emotion-based effects (EPN, LPP) or overall awareness effects (VAN, LP). Our primary focus was the investigation of connections between knowledge- and appearance-based emotional effects and visual awareness in the EPN component. We also considered the LPP as another component that might reflect enhanced processing of emotional information at a later (supposedly conscious) processing stage. To place emotion-specific effects in the context of overall processing during the attentional blink, we also report overall awareness differences (VAN and LP components). A more detailed examination of the time course of overall awareness-specific differences (as examined for the P1, N1, N2, and P3 components) for data from the same experiment has been reported in a separate article, which focused on the question of graded versus all-or-nothing awareness (Eiserbeck et al., 2022).

### 1.6. Pre-Registration

The hypotheses and methods of this study were pre-registered using the Open Science Framework (OSF) and can be accessed under https://osf.io/us754 (pre-registration 1; for the aspect of affective knowledge) and https://osf.io/2yspe (pre-registration 2; for the aspect of facial trustworthiness and its interaction with affective knowledge). Please note that in regard to the ERP analyses, we deviated from the pre-registration in a few points. In Appendix A2, we list and explain these changes.

## 2. Methods

The methods used in this study were largely based on those used in a previous behavioral study (Eiserbeck & Abdel Rahman, 2020) and extended to include the recording and analyses of event-related potentials. Data from the present experiment provided the basis for another article (Eiserbeck et al., 2022), which examined overall awareness-specific differences in the P1, N1, N2, and P3 components, focusing on the question of graded versus all-or-nothing awareness. In contrast, the present article is focused specifically on social-emotional influences on awareness.

### 2.1. Participants

The sample comprised 32 native German speakers (21 female; *M*_age_ = 26.1 years, *SD* = 6.65; normal or corrected-to-normal vision), in accordance with a pre-determined sample size based on power analyses, as described below. All participants provided written informed consent prior to participation. The study was conducted according to the principles expressed in the Declaration of Helsinki and was approved by the local Ethics Committee. Participants received either course credit or monetary compensation.

Based on pre-registered criteria, initial data sets of fifteen participants were discarded and replaced, keeping the defined number of thirty-two participants and ensuring a balanced within-subjects design. Data sets were replaced if one of the following criteria applied: T1-performance below 80% (8 participants); false alarm rate in T2-absent trials in the short lag condition above 50% (4 participants); failure to correctly recall the valence of the associated information for more than one-third of the 24 T2-faces, as assessed in a retrieval task at the end of the experiment (3 participants). These criteria were selected to ensure that person knowledge was learned sufficiently well and that enough trials for ERP analyses could be obtained without too many guess trials that would dilute the analyses.

Planning of the sample size was based on a behavioral pilot test (*N* = 5). We used a generalized linear mixed model predicting T2 awareness (hits defined by correct gender classification and at least a “strong impression” as the subjectively reported visibility) by affective knowledge (negative vs. neutral) and appearance (continuous predictor), including by-participant and by-item random intercepts. The resulting effect size for the interaction of affective knowledge and appearance (*b* = 0.16) was entered in an a priori power analysis in R with the SIMR package (Green & Macleod, 2016). We aimed for a power of at least 80% as conventionally deemed adequate (see Green & MacLeod, 2016). After running 1,000 randomizations given different sample sizes, results indicated that we would need to test 20 participants to detect an effect with an expected power of 83.90%, 95% CI [81.47, 86.13]. A further power analysis was run to estimate the sample size needed to detect a main effect of affective knowledge (*b* = 0.26), which yielded a similar result of 23 participants (expected power: 81%, 95% CI [78.4, 83.4]). For a balanced experimental design with a multiple of four participants and to have enough power to detect ERP effects, which may be smaller in size than the behavioral effects, we decided to test 32 participants.

### 2.2. Materials

#### 2.2.1. Pictures

T2 target stimuli consisted of 24 portraits of faces (12 female) with Caucasian appearance, displaying neutral emotional expressions, taken from the Chicago Face Database (CFD; Ma et al., 2015). Based on the rating data of the database, faces were chosen to cover a range of trustworthiness evaluations from low to high perceived trustworthiness. The pictures were converted to greyscale images and cropped so that no hair and ears were visible. The outer shape of the face was retained (instead of, e.g., applying an oval mask) because shape may be a factor affecting trustworthiness impressions (Kleisner et al., 2013). To minimize low-level confounds, histograms (i.e., the distributions of brightness values) of the images were equated using the SHINE toolbox (Willenbockel et al., 2010) in MATLAB R2016a.

Six additional faces (three female) from the CFD, processed the same way as the T2 faces, served as fillers associated with positive knowledge during learning. These additional faces served as positive anchor points in order to prevent biases that could arise due to only presenting neutral and negative knowledge. They were not presented in the attentional blink task.

To serve as distractor images in the attentional blink task, 12 additional faces (6 female) with average trustworthiness ratings were chosen from the database and processed in the same way as described above. Additionally, the facial features were cut out, rotated, and randomly placed in different positions within the face, thus creating abstract-looking faces. For each distractor, features of two faces of the same sex were “mixed” to further contribute to an abstract impression. Our aim was to create distractors that are visually similar to the T2 targets (see Müsch et al., 2012, for the importance of target-distractor-similarity) but sufficiently distinguishable.

T1 target stimuli consisted of 36 images displaying either the face of a dog or a similarly looking blueberry muffin, all converted to greyscale and cropped to the same oval shape.

Stimuli were presented on a 19-inch LCD monitor with a 75-Hz refresh rate. During all phases of the experiment, the images were displayed on a grey background with a size subtending 5.8° vertical visual angle and 4.3° horizontal visual angle (viewing distance: 70 cm).

#### 2.2.2. Person-Related Information

Twenty-four sentences describing negative or neutral social behavior were recorded by a male speaker in a neutral intonation (mean duration = 2.63 s). The sentences were rated in a web-based questionnaire (*N* = 20) on valence (negative: *M* = 1.67, *SD* = 0.36; neutral: *M* = 4.04, *SD* = 0.10; difference: *t*(11) = -31.3, *p* < .001), and arousal (negative: *M* = 4.94, *SD* = 0.54; neutral: *M* = 1.37, *SD* = 0.13); difference: *t*(11) = 22.2, *p* < .001), using a seven-point self-assessment manikin scale (Bradley & Lang, 1994). Six additional sentences describing a positive behavior served as fillers during learning (valence: *M* = 6.39, *SD* = 0.15; arousal: *M* = 4.58, *SD* = 0.33; valence difference to neutral sentences: *t*(7.45) = -33.99, *p* < .001; valence difference to negative sentences: *t*(15.85) = - 39.18, *p* < .001; arousal difference to neutral sentences: *t*(5.86) = -23.27, *p* < .001; no significant arousal difference to negative sentences: *t*(15.18) = 1.73, *p* = .104). Sentences always started with “she”/ “he” or “this woman”/ “this man,” followed by the description of a social behavior, e.g., “threatened a shop assistant with a knife” (negative knowledge condition) or “asked a waiter for the menu” (neutral knowledge condition). For a full list of sentences, see Appendix Table A1. The sentences were presented auditorily rather than visually since this enabled the simultaneous presentation of face stimulus and auditory information during the learning phase of the experiment without taking the visual focus away from the face. Furthermore, it allowed for a precise control of presentation times during the learning phase, without needing to consider differences in participants’ reading speed.

### 2.3. Procedure

A graphical overview of the different experimental phases can be found in Figure 1B.

#### 2.3.1. Learning Phase

##### 2.3.1.1. Pre-Learning Ratings of Trustworthiness and Facial Expression

Participants rated the trustworthiness and facial expression of all 30 faces (T2 target faces as well as filler faces associated with positive information in the learning phase) prior to knowledge acquisition. Ratings were completed block-wise with a counterbalanced order across participants. Faces were presented in random order within the blocks. At the beginning of each trial, a fixation cross was presented for 500 ms. Subsequently, a face was displayed for 1 s, followed by a short instruction and a 7-point scale. Depending on the task and stimulus gender, the instruction stated, “Please rate the trustworthiness of this woman[/man]” or “Please rate the facial expression of this woman[/man].” The ends of the scales were labeled, in case of the trustworthiness rating as “not at all trustworthy” (coded as 1) and “very trustworthy” (coded as 7) and for the facial expression rating as “negative” (coded as 1) and “positive” (coded as 7). The direction of the scales (left to right or right to left) was counterbalanced across participants. Participants used the left mouse button to indicate their choice. There were no time constraints for responses.

##### 2.3.1.2. Knowledge Acquisition

After completion of the ratings, participants acquired knowledge about the persons. To this end, each of the 30 faces was presented together with the accompanying auditory information. During each trial, first, a fixation cross was shown for 500 ms. Subsequently, the face was displayed for 6 s. Beginning at 1 s after face onset, the auditory information was presented via loudspeakers. Assignment of faces to affective knowledge conditions was counterbalanced across participants, such that each of the 24 T2-target faces was associated equally often with negative and neutral information. The filler faces were accompanied by the same information for all participants. To foster learning, each face was presented together with the accompanying information for a total of five times in blocks of gradually increasing numbers of faces (4, 6, or 12 faces from each affective knowledge condition plus 2, 3, or 6 filler faces) and simple judgment tasks related to the presented behaviors were included (e.g., “Is this person’s behavior common?”; Abdel Rahman, 2011; Baum et al., 2020; Suess et al., 2015). In this way, participants learned information for a few faces at a time through the help of repetition and answering of control questions, and the number of faces per block was then gradually increased. The rationale was to ensure that information for each face was learned to a sufficient degree (as later checked in a questionnaire at the end of the experiment). This design also ensured that each face was shown for the same amount of time during the learning phase.

After learning, the EEG was prepared and then recorded during the attentional blink task, the subsequent rating task, and an eye movement calibration procedure at the end of the experiment.

#### 2.3.2. Test Phase

##### 2.3.2.1. Attentional Blink Task

For each trial of the attentional blink task, first, a fixation cross was presented for 500 ms. Then, 13 pictures were shown in rapid succession (for illustration, see Fig. 1C), with a presentation time of 107 ms each and without a time interval between pictures. Regular trials contained 11 distractor images, presented in randomized order, and two targets: a dog or muffin (T1) and a face (T2). T2 (if present) was always presented as the 10^th^ stimulus, whereas T1 position varied: It was either presented as the 3^rd^ stimulus (entailing a lag of 7 items between T1 and T2; long lag) or as the 7^th^ stimulus (entailing a lag of 3 items; short lag). The task comprised 696 trials in total. As T1, in 50% of cases a dog was shown, and in 50% a muffin. All T2-faces were presented equally often—resulting in an equal number of trials for the two affective knowledge conditions (144 trials per affective knowledge condition for each short and long lag). To estimate the false alarm rate for each participant, T2 was absent in 60 trials (17%) within each lag; instead, another distractor was presented. All trial types (short or long lag, T2 present or absent, neutral, or negative knowledge) were presented in randomized order.

Participants were instructed to look for the dog/muffin and the face. They were informed that both targets are equally important but that not every sequence contains a face. After each trial, participants indicated via response keys (a) whether they saw the image of a dog or a muffin as T1 (options: *dog* / *muffin* / *I don’t know*), (b) whether they saw a male or a female face as T2 (options: *male* / *female* / *I don’t know*), and (c) how clear their subjective impression of T2 was on a four-point perception awareness scale (PAS; Ramsøy & Overgaard, 2004; options: *not seen* / *slight impression* / *strong impression* / *seen completely*). The response option “I don’t know” was included in both T1 and T2 tasks to keep participants from guessing since guess trials would dilute the ERP analyses of hit versus miss trials. There were no time constraints for answering.

Since the study was conducted in German and the original PAS labels do not translate well in a literal sense (in the authors’ subjective intuition as German speakers), slightly modified labels were used, representing German expressions with the closest meaning to the original labels (and to the more detailed explanation of each label provided with the PAS): “Nichts gesehen” (Engl.: not seen), “Leichter Eindruck” (Engl.: slight impression), “starker Eindruck” (Engl.: strong impression), and “vollständig gesehen” (Engl.: seen completely). Participants were instructed to select “not seen” if they had no impression at all of a human face being presented. They were further instructed that, if they had the impression of a face being presented, they should differentiate whether they had only a vague impression of the stimulus (“slight impression”), a fairly good but not full impression (“strong impression”) or whether they saw the stimulus completely (“seen completely”). They were encouraged to use the whole scale range. A previous study (and corresponding pre-study; Eiserbeck & Abdel Rahman, 2020) verified that the scale with the chosen labels provides a suitable measure of perceived visibility.

##### 2.3.2.2. Post-Learning Ratings of Trustworthiness and Facial Expression

The procedure of the second rating phase was identical to the first rating phase except that the tasks (trustworthiness and facial expression rating) were repeated three times. This was done to obtain enough trials for the ERP analyses. The analysis of ERPs during this post-learning rating phase served as an examination of knowledge and appearance-based effects under conditions of unimpeded perception. It was initially also planned as a localizer task for obtaining the regions and time intervals of interest for ERP analyses during the attentional blink task (but the strategy had to be changed ultimately; please see description in section 2.5.2.2).

After the experiment, the successful acquisition of person-related information was checked via a computerized survey. Participants indicated which kind of behavior (negative or neutral) was associated with each target face and what they recalled that particular behavior to be.

### 2.4. EEG Recording and Preprocessing

During the attentional blink task and post-learning ratings, the EEG was recorded with BrainAmpDC amplifiers, from Ag/AgCl electrodes (passive) at 62 scalp sites according to the extended 10–20 system (EasyCap M1 electrode cap). The sampling rate was 500 Hz and all electrodes were referenced to the left mastoid. An external electrode below the left eye was used to measure electrooculograms. During recording, low- and high-cut-off filters (0.016 Hz and 1000 Hz) were applied, and all electrode impedances were kept below 10 kΩ. After the experiment, a calibration procedure was used to obtain prototypical eye movements for later artifact correction with BESA (Ille et al., 2002), as described below. Processing and analyses of the data were based on the EEG-processing pipeline by Frömer et al. (2018). An offline pre-processing was conducted using MATLAB (Version R2016a) and the EEGLAB toolbox (Version 13.5.4b; Delorme & Makeig, 2004). The continuous EEG was re-referenced to a common average reference, and eye movement artifacts were removed using a spatiotemporal dipole modeling procedure with the BESA software (Ille et al., 2002). The corrected data were low-pass filtered at 40 Hz pass-band edge (zero-phase FIR-filter with transition band width of 10 Hz and cutoff frequency (-6 dB): 45 Hz). Subsequently, they were segmented into epochs of -200 to 1,000 ms relative to T2 onset and baseline-corrected using the 200 ms pre-stimulus interval. Segments containing artifacts (absolute amplitudes over ±150 µV or amplitudes changing by more than 50 µV between samples) were excluded from further analysis. In the rating task data, 511 of 5760 trials (8.87 %) were excluded based on artifacts (mean number of excluded trials per participant: 15.97 out of 180; *SD* = 15.46). The number of excluded trials per participant did not differ between the neutral (*M* = 9.38; *SD* = 9.53) and negative knowledge conditions (*M* = 9.56; *SD* = 9.19; *t*[31] = -0.30, *p* = .766). In the attentional blink task data, 1487 of 22272 trials (6.68 %) were excluded based on artifacts (mean number of excluded trials per participant: 46.47 out of 696; *SD* = 51.33). The number of excluded trials per participant did not differ between the neutral (*M* = 19.16; *SD* = 21.65) and negative knowledge conditions (*M* = 19.09; *SD* = 20.59; *t*[31] = 0.06, *p* = .953).

### 2.5. Data Analyses

#### 2.5.1. Behavioral Data

Behavioral data were analyzed using linear mixed models (LMM). Analyses were conducted in R (Version 4.2.2, R Core Team, 2018) using the lme4 package (Version 1.1-31; Bates et al., 2015) and the lmerTest package (Version 3.1-3; Kuznetsova et al., 2017) to calculate *p*-values via the Satterthwaite approximation in the case of linear mixed models. In the case of generalized linear mixed models (GLMM), *p*-values were based on the Wald *z*-test implemented in lme4. In all (G)LMM analyses, we aimed to include the maximal random effects structures justified by the design (Barr et al., 2013), with random effects for subjects and items. If models failed to converge, random effects were excluded based on least explained variance (using the *rePCA* function, see Bates et al., 2018). Facial appearance was treated as a continuous predictor, with the mean rating value of trustworthiness across participants before knowledge acquisition serving as consensus *appearance* score for each face.

##### 2.5.1.1. Manipulation check: Trustworthiness and Facial Expression Ratings

As manipulation checks, we examined evaluations during both rating phases. Rating data were analyzed with LMMs, with trustworthiness or expression rating serving as the dependent variable. The models included the fixed factors phase (before learning / after learning), affective knowledge (neutral/negative), and appearance. The predictors affective knowledge and appearance were nested within phase to specifically test effects before learning and after learning. Effect coding was applied for the factor affective knowledge (neutral: -0.5, negative: 0.5); the continuous predictor appearance was mean-centered.

##### 2.5.1.2. Main Task: Attentional Blink

Behavioral data of the attentional blink task were analyzed with binomial GLMMs, only including trials in which T1 was correctly identified to ensure that attention was paid to the first target as a pre-requisite for the attentional blink to occur. Hit (encoded as 1 for hit and 0 for miss) served as the dependent variable. As specified in the pre-registration, analyses were conducted separately for two criteria defining trials as T2 hit or miss (see Figure 4, Top). To count as a hit trial, for both criteria, the gender of T2 needed to be classified correctly. Furthermore, participants had to indicate either at least a *slight impression* (liberal hit criterion) or at least a *strong impression* (strict hit criterion) as the subjectively rated visibility of T2. These two different criteria were implemented since previous findings indicate that it may be important to take into account the threshold for considering a trial as hit or miss (see Eiserbeck & Abdel Rahman, 2020). Specifically, this takes into account that awareness may be a gradual phenomenon rather than a dichotomy between not seeing a stimulus and seeing a stimulus completely, and that boundaries could be drawn at different points on a continuum. While setting the boundary at a “slight impression” would mean that participants had at least a vague impression of the face as compared to not being aware of it at all, the boundary at a “strong impression” impression would mean that they perceived the stimulus fairly well as compared to having no or a vague impression. Effects of person knowledge were previously only observed for the later criterion (Eiserbeck & Abdel Rahman, 2020). To verify the presence of an (overall) attentional blink effect, GLMMs with the fixed factor lag (short/long) were computed. To test the hypotheses concerning affective knowledge and appearance-based trustworthiness, analyses were confined to short lag trials (as specified in the pre-registration), including the fixed factors affective knowledge (neutral/negative) and appearance. Effect coding was applied for the factors lag (short: -0.5, long: 0.5) and affective knowledge (neutral: -0.5, negative: 0.5); the continuous predictor appearance was mean-centered.

#### 2.5.2. ERP Analyses

##### 2.5.2.1. ERPs During Conscious Perception: Post-Rating Phase

We used cluster-based permutation (CBP) tests (Groppe et al., 2011; Maris & Oostenveld, 2007) to investigate EPN and LPP effects of affective knowledge, appearance, and their interaction during the second rating phase, i.e., under conditions of unimpeded conscious perception. This non-parametric approach enabled us to limit the time range and electrode sites to where and when EPN or LPP effects can be expected while at the same time allowing for variance regarding the location of effects and controlling for multiple testing. CBP tests were conducted in Matlab, using FieldTrip (Maris & Oostenveld, 2007), based on the implementation in Frömer et al. (2018). To specifically investigate EPN differences, the analysis was restricted to a time range and broadly defined posterior topographical region typical for the component (Abdel Rahman, 2011; Schacht & Sommer, 2009; Schupp et al., 2004; Suess et al., 2015), namely 150 to 350 ms after face stimulus onset including electrodes TP9, TP7, TP8, TP10, P7, P5, P3, Pz, P4, P6, P8, PO9, PO7, PO3, Poz, PO4, PO8, PO10, O1, Oz, and O2. Considering that the difference pattern observed in the EPN is sometimes found to begin already in the N170 (e.g., Baum & Abdel Rahman, 2021b; Luo et al., 2016), the time range of the N170 component (150-220 ms in the present data) is included as well. LPP effects were investigated within a typical time range and central topographical region (Abdel Rahman, 2011; Cuthbert et al., 2000; Schacht & Sommer, 2009; Schupp et al., 2004), from 400 to 800 ms including electrodes FC3, FC1, FC2, FC4, C3, C1, Cz, C2, C4, CP3, CP1, CPz, CP2, CP4, P3, Pz, P4, PO3, POz, and PO4. For transparency, we would like to note that these electrodes and time ranges were not pre-registered in this specificity but chosen on the basis of previous studies as indicated above. Since CBP tests in the current implementation are based on the comparison of two conditions, the continuous variable appearance was converted into a factor with two levels by using a median split to separate between low- and high-trustworthy-looking faces. To investigate interactions between affective knowledge and appearance, we used a double subtraction procedure, comparing the difference between negative and neutral knowledge in the untrustworthy and trustworthy appearance condition ([Untrustworthy: Negative-Neutral] – [Trustworthy: Negative-Neutral]). A dependent samples t-statistic was used to evaluate the effect at the sample level to determine cluster inclusion, using an alpha level of .05 for a single test. To be included in a cluster, a minimum number of two significant neighborhood channels was required. Thereby, an electrode’s spatial neighborhood was defined as adjacent electrodes in the cap. As the test statistic on the cluster level, the maximum of the cluster-level statistics was used (i.e., the largest sum of sample-specific t-statistics for each of the different clusters produces the test statistic), and a two-tailed test was applied. The number of permutations was 10,000.

##### 2.5.2.2. Main Task: Attentional Blink

###### Planned Analyses

We had originally planned to use the post-learning rating phase as a localizer task for ERP effects and then analyze the same regions of interest and time range in the attentional blink task to examine the presence of ERP effects after T2-face presentation during short lag attentional blink trials. This approach was unsuccessful, likely due to task differences and interactions with awareness. We report details and results of this analysis in Appendix A2.2.

###### Exploratory Analyses

Considering that effects of emotional information may be temporally and topographically shifted in the attentional blink task compared to the rating task due to task differences and that they might be observable only in interaction with awareness, we further investigated differences using CBP tests. This approach enabled us to examine effects within the EPN/VAN and LPP/LP time ranges and regions of interest while allowing for variance regarding the exact timing and location of effects and controlling for multiple testing. Settings for the CBP tests were the same as reported above. Due to the overlaps in the typical latency and topography between EPN and VAN, and LPP and LP, respectively, within each set of components (EPN/VAN and LPP/LP), CBP tests were based on the same broad region and time interval (as described for EPN and LPP above). We examined main effects of knowledge, appearance and the interaction of knowledge and appearance, as well as main effects of awareness. Furthermore, and most relevant to the research question, we examined interactions between knowledge and awareness (implemented as double difference: [Negative: Hit-Miss]-[Neutral: Hit-Miss]), as well as appearance and awareness (implemented as double difference: [Untrustworthy: Hit-Miss] – [Trustworthy: Hit-Miss]). Analogous to the behavioral data analysis, only short lag trials in which T2 was present and T1 was correctly identified were examined. Likewise, we again examined both hit criteria (hits defined as trials with correct classification and either an at least slight or strong impression as subjectively rated visibility). Different from the behavioral analyses, we defined as misses only those trials in which participants chose the response option “I don’t know” in the T2 classification task combined with a visibility rating not higher than “not seen” (liberal criterion) or than “slight impression” (strict criterion). Other trials (e.g., wrong answers or low visibility ratings combined with correct gender classification) were excluded from CBP analyses. This was due to the following considerations: Generally, we opted to include as many trials as possible and kept the criteria for the behavioral analyses analogous to those used in a previous study (Eiserbeck & Abdel Rahman, 2020) for comparability. In CBP tests of the EEG data, however, we chose to restrict the analyses to those miss trials which represent the clearest distinction from the hit trials, in order to clearly localize the relevant ERP differences associated with visual awareness of a face.

Significant interactions in CBP tests were followed up with linear mixed models predicting mean cluster activity by the respective conditions to investigate the specific directions of effects. To this end, based on the topographical and temporal distribution of the observed cluster, single-trial mean amplitudes were obtained and entered as the dependent variable in the models. Unlike the CBP tests, this allowed us to control for by-participant and by-item random variance.

Finally, to predict awareness (hit/miss) based on all relevant variables and their interaction within one model, we conducted GLMM analyses. This served as an additional validation and further examination of an observed interaction between knowledge and awareness in the CBP tests. It also enabled examining a potential three-way interaction with appearance, which would have been overly complex to include in CBP tests. Since neural activity precedes behavior and we aimed to predict awareness by neural activity, in these models, hits were predicted by mean amplitudes rather than the other way around. Based on the topographical and temporal distribution of the observed cluster, single-trial mean amplitudes were obtained and mean-centered. We extended the previously specified GLMM described in the behavioral analyses by the predictor *mean cluster amplitude*: Hits (0/1) were predicted by knowledge (neutral/negative), appearance (continuous predictor), and mean cluster amplitude (continuous predictor), including all interactions between the predictors, as well as the previously specified random effects structure

## 3. Results

### 3.1. Ratings and Manipulation Checks

#### 3.1.1. Behavioral Data

Figure 2 provides an overview of the ratings before and after knowledge acquisition. A highly similar pattern of results was observed for trustworthiness and facial expression ratings: Before learning, there was a main effect of appearance (trustworthiness: *b* = 1.00, *t*[41.08] = 8.47, *p* < .001; expression: *b* = 0.92, *t*[42.88] = 6.43, *p* < .001), with more positive ratings for more trustworthy compared to less trustworthy faces (i.e., mean appearance scores—representing “consensus” tendencies across all participants—predicted individual participants’ trustworthiness and expression ratings). Ratings for faces associated with neutral and negative knowledge did not differ significantly before knowledge acquisition (trustworthiness: *b* = -0.06, *t*[51.19] = -0.56, *p* = .577; expression: *b* = -0.07, *t*[62.50] = -0.76, *p* = .453), and there was no interaction between appearance and affective knowledge (trustworthiness: *b* = 0.07, *t*[56.64] = 0.51, *p* = .610; expression: *b* = 0.02, *t*[173.09] = 0.13, *p* = .895). After learning, appearance still predicted trustworthiness and expression ratings (trustworthiness: *b* = 0.70, *t*[41.08] = 5.93, *p* < .001; expression: *b* = 0.80, *t*[42.88] = 5.64, *p* < .001). Furthermore, there was a main effect of affective knowledge in both trustworthiness and expression task: Faces associated with negative information were rated as less trustworthy (*M* = 3.29) than faces associated with neutral information (*M* = 4.32; *b* = -1.00, *t*[51.19] = -8.72, *p* < .001), and the expressions of faces associated with negative information were rated as more negative (*M* = 3.70) than the expressions of faces associated with neutral information (*M* = 4.13; *b* = -0.42, *t*[62.50] = -4.69, *p* < .001). There was no significant interaction between appearance and affective knowledge (trustworthiness: *b* = 0.13, *t*[56.64] = 0.96, *p* = .340; expression: *b* = 0.15, *t*[173.09] = 1.25, *p* = .215).

**Figure 2.**
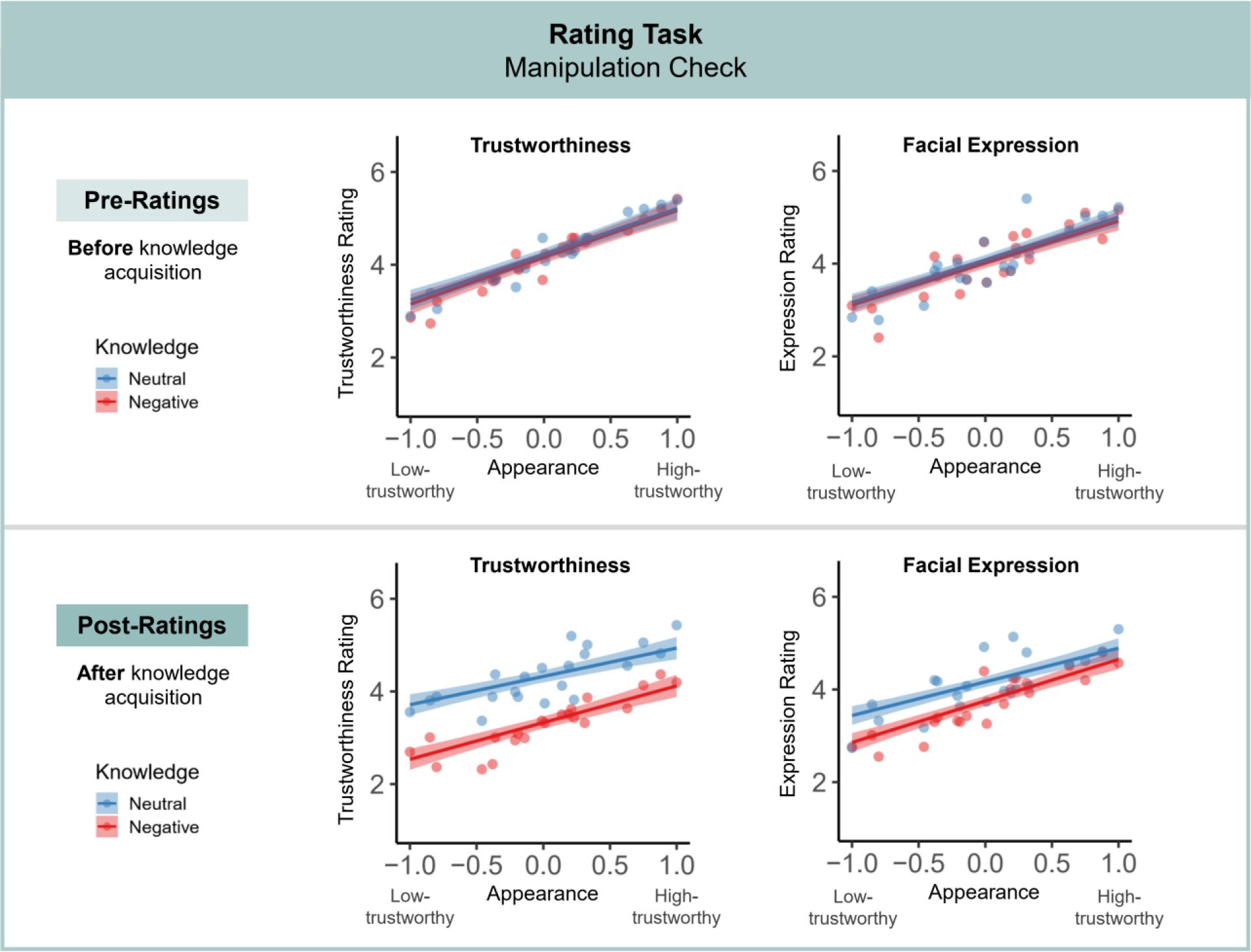
Manipulation check. Trustworthiness and facial expression ratings show a successful manipulation of affective knowledge and appearance: Before learning (top row), only appearance influenced ratings; after learning, there were additive effects of both affective knowledge and appearance (bottom row). The ratings were given on 7-point scale with the ends of the scales labeled—in case of the trustworthiness rating as “not at all trustworthy” (1) and “very trustworthy” (7) and for the facial expression rating as “negative” (1) and “positive” (7). The regression lines show partial effects from the LMM analyses; error bands depict ± 1 standard error; dots illustrate (descriptive) mean ratings for individual faces.

#### 3.1.2. ERPs During Conscious Perception: Second Rating Phase

In the EPN time range, CBP tests revealed a significant main effect of knowledge (maximal cluster: sum(*t*) = -184.59; *p* = .048), with amplitude values for faces associated with negative compared to neutral knowledge being significantly more negative (see Figure 3). The corresponding negative cluster was observed between 270 and 288 ms at electrodes P4, P6, P8, PO3, POz, PO4, PO8, PO10, Oz, and O2. No significant differences were found for appearance (maximal cluster: sum(*t*) = -93.03; *p* = .134) or the interaction of knowledge and appearance (maximal cluster: sum(*t*) = 33.04; *p* = .279). In the LPP time range, no significant differences were found for either knowledge (maximal cluster: sum(*t*) = 33.04; *p* = .057), appearance (maximal cluster: sum(*t*) = -87.16, *p* = .174), or an interaction of knowledge and appearance (maximal cluster: sum(*t*) = 119.34, *p* = .140).

**Figure 3.**
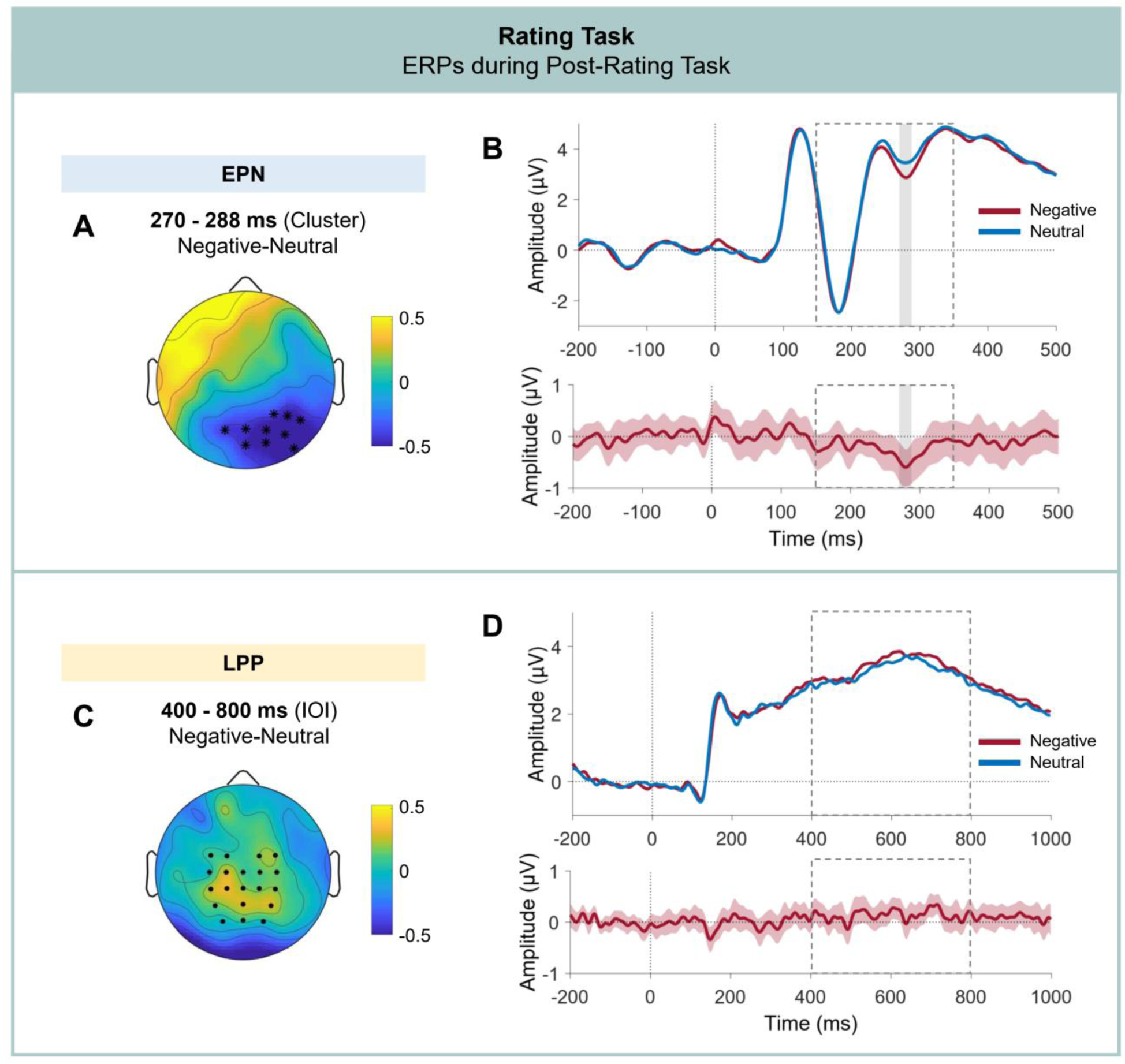
ERP differences under conditions of unimpeded conscious perception in the post-rating phase. Cluster-based permutation tests revealed a significant difference between faces associated with negative and neutral knowledge in the EPN, with enhanced negative amplitudes in the negative knowledge condition. In the LPP, no significant differences were found. (A) Difference topography for the affective knowledge effect in the EPN cluster interval. Asterisks mark the electrodes included in the cluster. (B) Grand-average event-related potential for each knowledge condition, based on pooled activity across electrode sites included in the cluster (upper panel), and corresponding difference curve with 95% bootstrap confidence interval (lower panel). The dashed frame indicates the time range included in the CBP test; the gray frame indicates the temporal extent of the cluster. (C) Difference topography for the negative versus neutral knowledge condition in the interval of interest (IOI) included in the CBP test. Dots mark the electrodes included in the CBP test. (D) Grand-average event-related potential for each knowledge condition, based on pooled activity across electrode sites included in the CBP test (upper panel), and corresponding difference curve with 95% bootstrap confidence interval (lower panel). The dashed frame indicates the time range included in the CBP test.

### 3.2. Main Task: Attentional Blink

#### 3.2.1. Behavioral Data

The mean T1 recognition rate was 91.06% (CI ± 0.38). The mean correct rejection rate in T2 absent trials was 85.37% (CI ± 1.12). Binomial GLMM analyses predicting hits/misses (1/0) by lag (short/long) confirmed the presence of an attentional blink effect, i.e., an effect of lag, for both liberal (*b* = 0.95, *z* = 5.77, *p* < .001) and strict criterion (*b* = 0.76, *z* = 4.53, *p* < .001), with higher hit rates in the long as compared to the short lag condition (liberal criterion: 83.32%, CI ± 1.09 vs. 69.52%, CI ± 1.25; strict criterion: 48.64%, CI ± 1.25 vs. 36.74%, CI ± 1.22).

Table 1 contains estimates (regression coefficients *b*) of the fixed effects, standard errors, and *z*- and *p*-values for the analyses of short lag trials for both hit criteria. Graphical illustrations of hit rates (i.e., the proportion of T2 hits in T1-correct trials) and distributions can be found in Figure 4. For the liberal criterion (hit = correct gender classification and at least visibility rating 2, “slight impression”; miss = all other trials), neither the main effects of affective knowledge or appearance nor their interaction reached statistical significance. In contrast, for the strict criterion (hit = correct gender classification and at least visibility rating 3, “strong impression”; miss = all other trials), a main effect of affective knowledge was found. Mean hit rates were higher in the negative (38.23%, CI ± 1.42) relative to the neutral knowledge condition (35.27%, CI ± 1.39). The main effect of appearance (*p* = .082) and the interaction effect of affective knowledge and appearance (*p* = .102) did not reach statistical significance.

**Table 1:** GLMM Statistics for Analysis of Short Lag Trials.

| Variable | Liberal hit criterion |  |  |  | Strict hit criterion |  |  |  |
| --- | --- | --- | --- | --- | --- | --- | --- | --- |
| | $b$ | $SE$ | $z$ | $p$ | $b$ | $SE$ | $z$ | $p$ |
| Intercept | 0.99 | 0.20 | 4.87 | **<.001** | -1.12 | 0.38 | -2.96 | **.003** |
| Knowledge (Neg-Neu) | 0.01 | 0.07 | 0.13 | .897 | 0.21 | 0.09 | 2.25 | **.025** |
| Appearance | -0.19 | 0.12 | -1.60 | .111 | -0.29 | 0.17 | -1.74 | .082 |
| Knowledge:Appearance | 0.09 | 0.11 | 0.78 | .437 | 0.23 | 0.14 | 1.64 | .102 |
| Random effects | Var. | SD |  |  | Var. | SD |  |  |
| Participants (Intercept) | 1.20 | 1.10 |  |  | 4.16 | 2.04 |  |  |
| Knowledge | 0.02 | 0.15 |  |  | 0.04 | 0.21 |  |  |
| Appearance | 0.12 | 0.34 |  |  | 0.11 | 0.34 |  |  |
| Knowledge:Appearance | 0.02 | 0.13 |  |  | 0.06 | 0.25 |  |  |
| Items (Intercept) | 0.08 | 0.28 |  |  | 0.20 | 0.45 |  |  |
| Knowledge | 0.03 | 0.18 |  |  | 0.05 | 0.21 |  |  |

*Note.* Neg = negative, Neu = neutral; higher *Appearance* value corresponds to a more trustworthy looking appearance; “:” indicates interactions between fixed factors.

**Figure 4.**
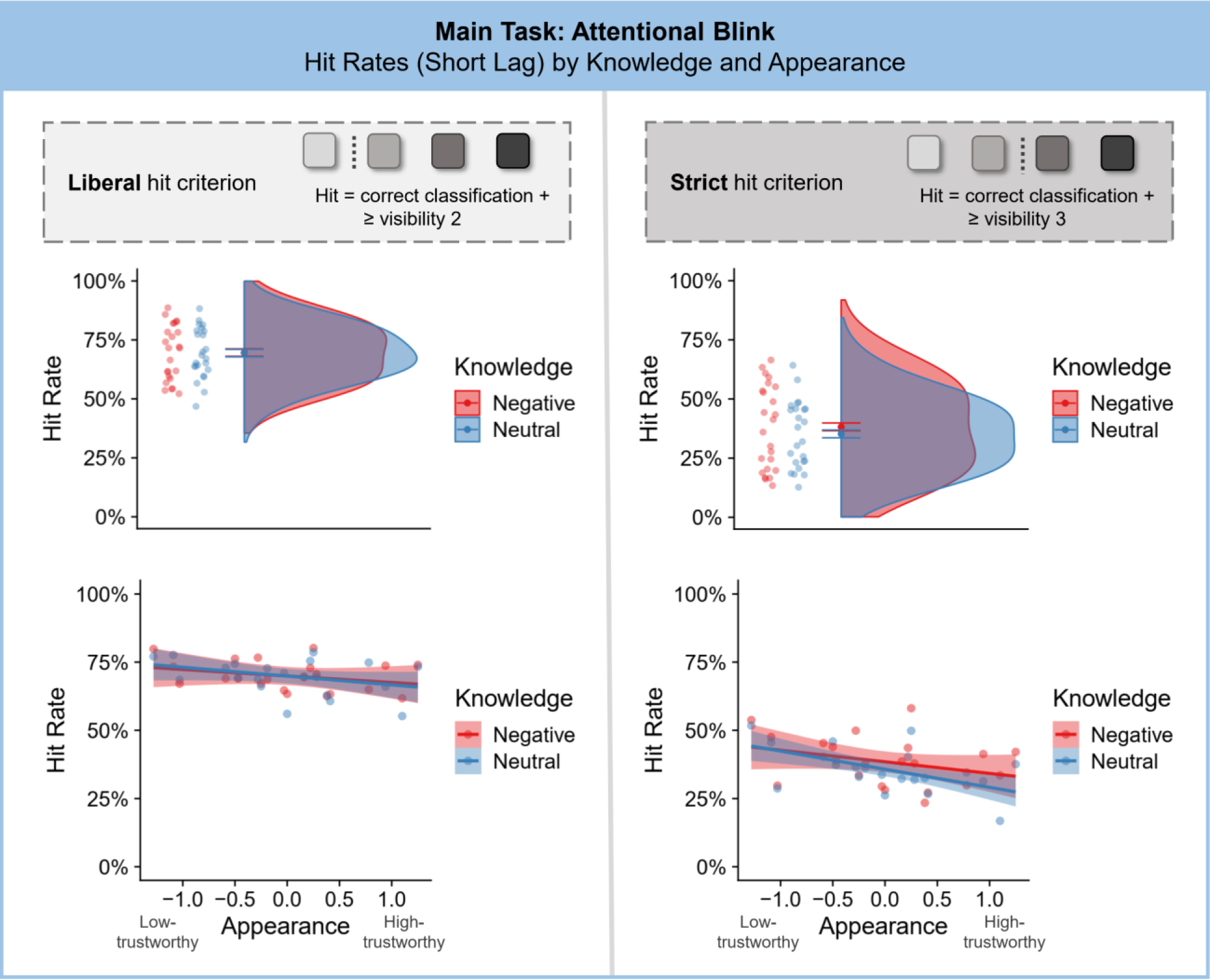
Behavioral results of the attentional blink task. Top: Overview of the two hit criteria implemented in the analyses, representing two differently stringent thresholds for considering a trial as a T2 hit or miss. Middle: By-item hit rates (ratio of T2 hits in T1-correct trials) and distributions in short lag T2-present trials depending on knowledge condition for liberal and strict hit criterion. Error bars depict 95% confidence intervals. Analyses on a single trial basis using GLMMs revealed a significant main effect of affective knowledge for the prediction of hit or miss for the strict but not for the liberal hit criterion. Bottom: Hit rates based on appearance scores and knowledge (with inter-subject variance controlled), with regression lines for appearance within each knowledge condition. Error bands depict ± 1 standard error.

#### 3.2.2. ERP Results (Exploratory Analyses)

##### 3.2.2.1 Trial numbers

Because it depended on participants’ performance in the attentional blink, the number of trials taken into account for each condition in the awareness-based analyses varied between participants. Per participant, an average number of 49 trials (SD = 37) per individual knowledge×awareness or appearance×awareness condition entered the analyses. Most participants achieved an acceptable number of trials (>10) in all conditions. However, 14 out of 32 participants had fewer than 10 trials in at least one of the individual conditions; 9 of those participants even had less than 5 trials in at least one condition. This was primarily due to a low number of miss trials due to high task performance or a low number of hit trials based on the strict criterion (i.e., rarely choosing the visibility options “strong impression” or “seen completely”). We had not pre-defined a minimum number of trials necessary in any one condition as an exclusion criterion. (This is a limitation of the present study.) Since it might introduce selection biases in the sample (and there already were stringent selection criteria in place, see section 2.1), we also chose not to replace data sets of participants a posteriori. Thus, the below reported analyses include data of all 32 participants. In the specific case of the strict hit criterion, 3 participants did not contribute data to the CBP tests, as they had 0 strict hit trials. To assess the impact of low trial numbers on the results, we conducted additional control analyses for the effects most relevant to the conclusions of the present article (interaction of knowledge and awareness in the EPN/VAN component), excluding those participants with less than 10 trials in any one condition (see Appendix A4). These analyses yielded highly similar results compared to the analyses including all participants reported below, although the differences failed to cross the threshold of statistical significance for the strict hit criterion in this case (*p* = .064; descriptive differences indicated the same direction). ERP curves and topographies corresponding to these control analyses (see Appendix A4, Figure A4.1) also show a highly similar pattern compared to those of the main analyses (see Figure 5). This indicates that it is unlikely that the respective results of the main analyses are due to noisy outliers.

**Figure 5.**
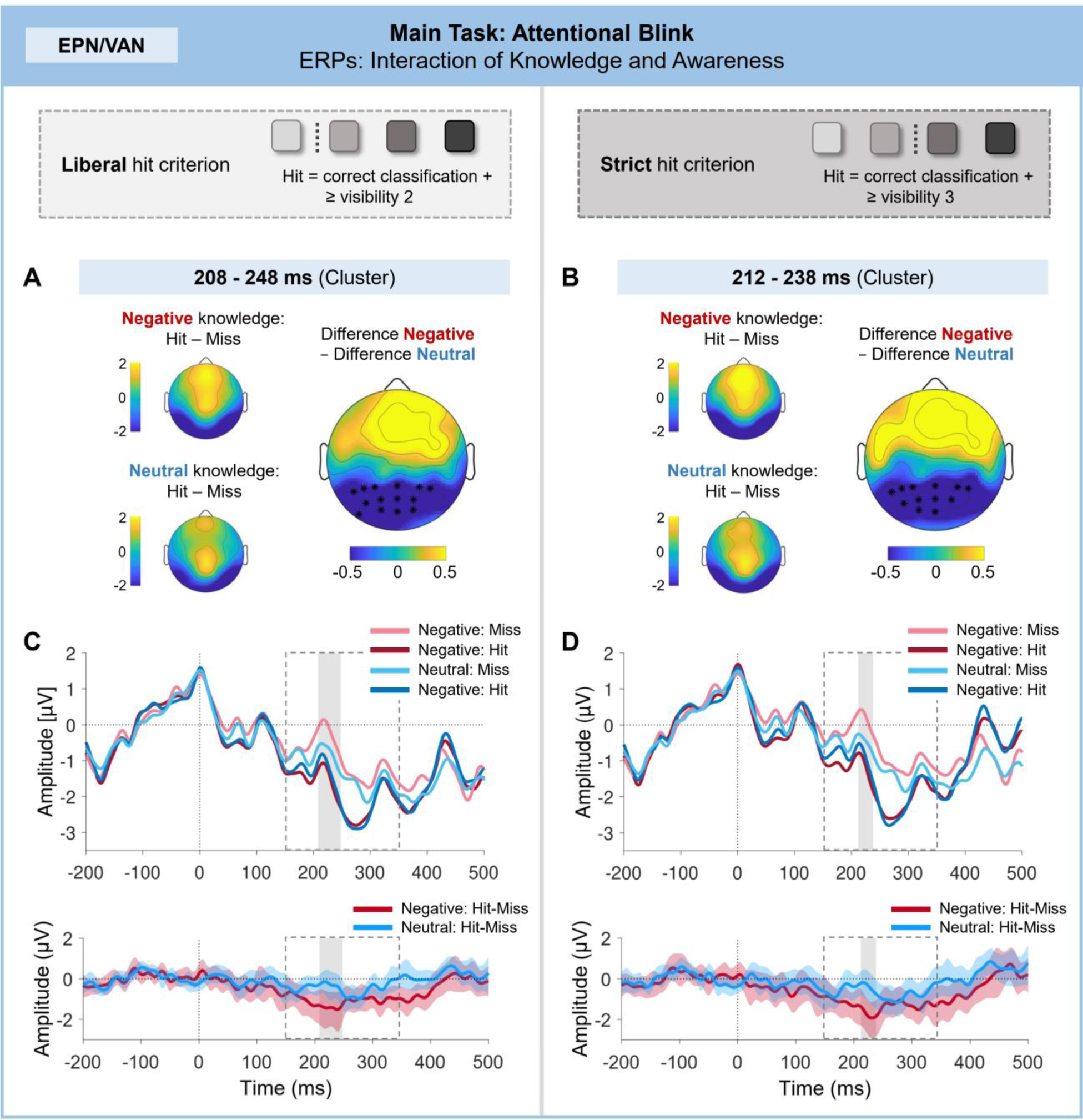
ERP differences during the EPN/VAN time range in the attentional blink: Interaction of knowledge and awareness. Cluster-based permutation tests revealed a significant interaction of knowledge and awareness in the EPN time range for both the liberal and strict hit criterion. (A), (B) The smaller topographies on the left show the difference between Hit and Miss trials in the negative and neutral knowledge conditions in the interval of the cluster. The topography on the right shows the corresponding double difference (awareness difference in negative condition – awareness difference in neutral condition). Asterisks mark the electrodes included in the cluster. (C), (D) Grand-average event-related potential for each knowledge × awareness condition, based on pooled activity across electrode sites included in the cluster (upper panel), and corresponding difference curves with 95% bootstrap confidence intervals for Hit-Miss trials in the negative and neutral knowledge conditions (lower panel). The dashed frame indicates the time range included in the CBP test; the gray frame indicates the temporal extent of the cluster.

**Figure 6.**
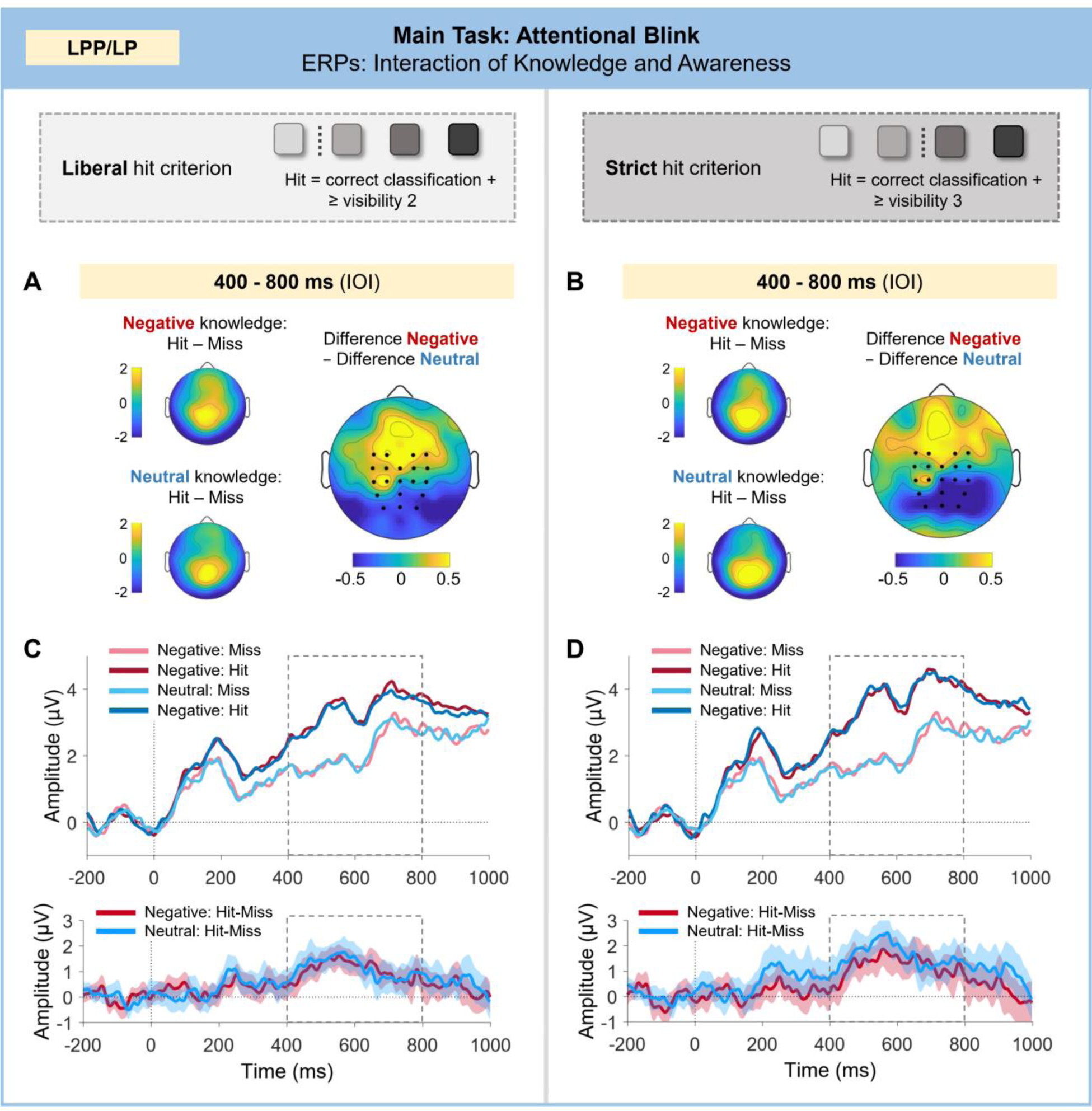
ERP differences during the LPP/LP time range in the attentional blink. Cluster-based permutation tests did not indicate an interaction of knowledge and awareness in the LPP/LP time range. (A), (B) The smaller topographies on the left show the difference between Hit and Miss trials in the negative and neutral knowledge conditions in the interval of interest (IOI) included in the CBP test. The topography on the right shows the corresponding double difference (awareness difference in negative condition – awareness difference in neutral condition). Dots mark the electrodes included in the CBP test. (C), (D) Grand-average event-related potential for each knowledge × awareness condition, based on pooled activity electrode sites included in the CBP test (upper panel), and corresponding difference curves with 95% bootstrap confidence intervals for Hit-Miss trials in the negative and neutral knowledge conditions (lower panel). The dashed frame indicates the time range included in the CBP test.

##### 3.2.2.2 Main and interaction effects of knowledge and appearance

In the EPN/VAN time range, CBPTs over all valid short lag trials regardless of awareness, revealed a significant difference between low-trustworthy and high-trustworthy looking faces (maximal cluster: sum(*t*) = 278.72, *p* = .020). The corresponding positive cluster was observed between 180 and 206 ms at electrodes P5, P3, Pz, P4, P6, PO7, PO3, POz, PO4, PO8, O1, Oz, and O2 (see Figure A5.1 in the Appendix). No main effect of affective knowledge (maximal cluster: sum(*t*) = 6.23, *p* = .359), or interaction of affective knowledge and appearance (maximal cluster: sum(*t*) = -107.56, *p* = .131) was observed.

In the LPP/LP time range, no significant differences were observed (main effect of affective knowledge: maximal cluster: sum(*t*) = 109.40, *p* = .166; main effect of appearance: maximal cluster: sum(*t*) = 268.42, *p* = .063; interaction of affective knowledge and appearance: maximal cluster: sum(*t*) = 235.27, *p* = .067).

##### 3.2.2.3 Main effects of awareness

In the EPN/VAN time range, CBP tests revealed a significant difference between hit and miss trials for both liberal hit criterion (maximal cluster: sum(*t*) = -5610.79, *p* = <.001) and strict hit criterion (maximal cluster: sum(*t*) = -6415.16, *p* = <.001), with more negative amplitude values for hit compared to miss trials. For both hit criteria, a similar, large negative cluster ranging across the whole tested time interval was observed (150-350ms) including almost all of the tested electrodes (both criteria: TP9, TP7, TP8, TP10, P7, P5, P3, P4, P6, P8, PO9, PO7, PO3, PO4, PO8, PO10, O1, Oz, and O2; strict criterion additional contained POz).

In the LPP/LP time range, CBP tests revealed a significant difference between hit and miss trials for both liberal hit criterion (maximal cluster: sum(*t*) = 11233.87, *p* < .001) and strict hit criterion (maximal cluster: sum(*t*) = 12508.99, *p* < .001), with more positive amplitude values for hit compared to miss trials. For both hit criteria, a similar, large positive cluster ranging across the whole tested time interval was observed (400-800ms) including all of the tested electrodes (both criteria: FC3, FC1, FC2, FC4, C3, C1, Cz, C2, C4, CP3, CP1, CPz, CP2, CP4, P3, Pz, P4, PO3, POz, and PO4).

A graphical illustration of these overall awareness differences in VAN and LP can be found in Figures A5.2 and A5.3 in the Appendix.

##### 3.2.2.4 Interactions of awareness with knowledge and appearance

In the EPN/VAN time range, CBP tests revealed a significant interaction of awareness and affective knowledge for both the liberal (maximal cluster: sum(*t*) = -474.08; *p* = .011) and strict hit criterion (maximal cluster: sum(*t*) = -285.50; *p* = .023), indicating an enhanced negative difference between hit and miss trials in the negative as compared to the neutral knowledge condition (see Figure 5). A corresponding negative cluster showed a similar distribution for both liberal and strict hit criterion: The cluster for the liberal hit criterion was observed from 208 to 248 ms at electrodes P7, P5, P3, Pz, P4, P6, PO9, PO7, PO3, POz, PO4, O1, Oz, and O2. The cluster for the strict hit criterion was observed from 212 to 238 ms at electrodes P7, P5, P3, Pz, P4, P6, PO7, PO3, POz, PO4, O1, and Oz. No interaction between appearance and awareness was observed in the CBP tests (no clusters). In the LPP/LP time range, no significant differences were observed for any of the comparisons (knowledge×awareness, liberal criterion: maximal cluster: sum(*t*) = 47.99, *p* = .291; knowledge×awareness, strict criterion: maximal cluster: sum(*t*) = -18.20, *p* = .373; appearance×awareness, liberal criterion: no clusters; appearance×awareness, strict criterion: no clusters).

We followed up on the observed significant interaction of awareness and affective knowledge with LMMs. To this end, single-trial mean amplitudes were obtained based on the topographical and temporal distribution of the corresponding cluster for the awareness × knowledge interaction—approximately 210 to 240 ms at electrodes P7, P5, P3, Pz, P4, P6, PO7, PO3, POz, PO4, O1, and Oz (corresponding to a rounded time range and electrodes included in the cluster for both hit criteria). These cluster amplitudes served as the dependent variable in the models. We examined effects of awareness on cluster amplitudes separately in the negative and neutral knowledge conditions. Significant effects of awareness were found in the negative condition, with more negative amplitude values in hit compared to miss trials (liberal criterion: *b* = -1.03, *t*[3639.65] = -5.00, *p* < .001; strict criterion: *b* = -1.57, *t*[1809.53] = -6.18, *p* < .001). In contrast, no significant differences were observed in the neutral condition (liberal criterion: *b* = -0.05, *t*[3666.46] = -0.25, *p* = .805; strict criterion: *b* = -0.35, *t*[1849.70] = -1.45, *p* = .148).

To further investigate the pattern of results, we also considered another direction of follow-up tests with LMMs, examining effects of knowledge on cluster amplitudes separately in hit and miss trials. No significant effect of affective knowledge on cluster amplitudes during hit trials was observed (liberal criterion: *b* = -0.19, *t*[5126.56] = -1.43, *p* = .152; strict criterion: *b* = -0.17, *t*[2888.81] = -0.93, *p* = .350). In miss trials, on the other hand, a significant main effect of affective knowledge was found, with more positive amplitude values in the negative as compared to the neutral condition (liberal criterion: *b* = 0.56, *t*[1126.16] = 2.71, *p* = .007; strict criterion: *b* = 0.61, *t*[1157.86] = 3.06, *p* = .002).

Finally, to predict awareness (hit/miss) based on all relevant variables and their interaction within one model, we conducted GLMM analyses. Thereby, we extended the previously specified GLMM described in the behavioral analyses by the predictor *mean cluster amplitude*: Hits (0/1) were predicted by knowledge (neutral/negative), appearance (continuous predictor) and mean cluster amplitude (mean-centered, continuous predictor), including all interactions between the predictors, as well as the random effects structure previously specified in the behavioral analyses. The models revealed a significant interaction between affective knowledge and mean amplitude for the liberal hit criterion (*b* = -0.04, *z* = -3.71, *p* < .001) as well as for the strict hit criterion (*b* = -0.03, *z* = -2.28, *p* = .023). To follow up on this significant interaction, we computed nested models in which the simple effects of mean cluster amplitude and appearance on hits are examined within each knowledge condition. The results of these nested models showed that the mean amplitude only has predictive value, or a higher predictive value respectively, for determining a trial as a hit or miss in the negative knowledge condition (liberal criterion: *b* = -0.05, *z* = -5.92, *p* < .001; strict criterion: *b* = -0.05, *z* = - 5.43, *p* < .001) as compared to the neutral knowledge condition (liberal criterion: *b* = -0.01, *z* = -0.71, *p* = .476; strict criterion: *b* = -0.02, *z* = -2.17, *p* = .030). For full model output, see Appendix Tables A3.1 and A3.2.

## 4. Discussion

Can social-affective knowledge about persons affect the access of faces to visual consciousness? In the present study, we investigated this question while additionally considering the impact of facial trustworthiness as a source of affective information derived from visual appearance. As a manipulation check, we tested and observed effects of affective knowledge and facial appearance in explicit evaluations of trustworthiness and emotional expressions. Knowledge- and appearance-based effects did not interact (see Fig. 2). In the attentional blink, replicating previous behavioral findings (Eiserbeck & Abdel Rahman, 2020), stimulus awareness under conditions of reduced attention was affected by affective knowledge about the person and not by their appearance (see Fig. 4). This influence of knowledge on visual awareness was also reflected in the ERPs (see Fig. 5): Cluster-based permutation tests revealed an effect in the time range of the early posterior negativity (EPN)—a component associated with enhanced attention and facilitated processing of emotional stimuli. Specifically, in the negative knowledge condition compared to the neutral condition, activity during this time range had an enhanced predictive value in determining the awareness of faces. No clear evidence for an impact of facial trustworthiness in the attentional blink was found.

### 4.1. Manipulation Check: Evaluations of Trustworthiness and Facial Expressions

The trustworthiness and facial expression evaluations after learning demonstrate successful manipulations of affective knowledge and appearance. In line with previously reported effects of affective person knowledge, faces associated with negative information were rated as less trustworthy, and their emotional expressions were rated as more negative than those of faces associated with relatively neutral knowledge (Eiserbeck & Abdel Rahman, 2020; Suess et al., 2015; Verosky et al., 2018). The consensus trustworthiness score across participants for each stimulus predicted individual trustworthiness and expression ratings before as well as after learning of affective information. This shows the systematic agreement between participants in regard to trustworthiness inferences from appearance and the strong connection of trustworthiness judgments to facial expression ratings, in line with the emotion-overgeneralization account of facial trustworthiness (Zebrowitz & Montepare, 2008). Notably, even with verbal information about a person’s previous behavior available, faces with a less trustworthy appearance (as based on consensus ratings) still received lower trustworthiness ratings compared to more trustworthy looking faces. This demonstrates the persistence of trustworthiness inferences based on appearance, even in the presence of more explicit (and potentially more diagnostic) social information, which is in line with previous findings (Jaeger et al., 2019). No interaction between affective knowledge and appearance was observed.

### 4.2. ERPs during Unimpeded Perception (Post-Learning Evaluations)

In the ERPs corresponding to unimpeded conscious face perception during post-learning evaluations, systematic differences in brain activity were observed for affective knowledge but not for appearance or an interaction of both factors. For affective knowledge, an effect with a pattern typical for the EPN component was found, with enhanced negative mean amplitudes for faces associated with negative as compared to neutral knowledge. The direction of the effect as well as the topographical and temporal distribution of the observed cluster—posterior sites between 270 to 288 ms after stimulus onset—is in line with previous evidence (Abdel Rahman, 2011; Luo et al., 2016; Suess et al., 2015; Wieser et al., 2014; Xu et al., 2016). Regarding the relatively narrow time range of the observed cluster, the following aspects have to be considered: First, while the result of the CBP analysis demonstrates a significant effect of affective knowledge within the tested latency range of 150 to 350 ms and posterior electrodes, the temporal and topographical extent of the cluster cannot be interpreted in terms of the actual extent of the effect, which may be larger in size than the single adjacent electrode×time points that crossed the threshold of statistical significance (Maris & Oostenveld, 2007; Sassenhagen & Draschkow, 2019). Second, the post-learning rating phase took place at the very end of the experiment, after the extensive attentional blink task, which represented the main focus of the present study. The temporal distance from the learning phase as well as decreasing mental focus towards the end of the experiment may have led to a reduction in the size of the EPN effect. No LPP effect was observed for affective knowledge. This might be due to the fact that in the evaluation task, participants were asked for spontaneous impressions of trustworthiness and expression, which do not necessitate an intentional retrieval of or focus on the associated emotional information. Consequently, differences are found in the EPN component as a marker of early reflexive emotional processing and not in the LPP component, typically indicative of later, more elaborate emotional processing. This consideration is in line with the finding that task relevance modulates emotion effects during the LPP, whereas emotion effects during the EPN are independent of task relevance (Schindler & Straube, 2020).

Contrary to expectations, no EPN or LPP effects were found for appearance. This might be due to the following reasons: We used CBP tests to examine the presence of ERP effects, which enabled us to restrict the time range and electrode sites to where effects are expected while not necessitating a precise a priori definition of the distribution of the effects, which often vary between studies for these components. Due to the correction for multiple testing, this approach is more conservative than, e.g., an analysis of mean amplitudes in a time range obtained through visual inspection, and it prevents the problem of circular analysis (that is, defining the latency or topography of the effect based on the data itself and then using it for analysis, see Luck & Gaspelin, 2017). A caveat of this approach is that it is harder to detect effects of small temporal and topographical distribution, especially for components with a less central topographical distribution (since electrodes at the outer end of the electrode cap possess fewer neighboring electrodes, reducing the chances for building a cluster). Even though visual inspection indicated differences in the expected direction in the EPN time range with enhanced negative amplitudes for less trustworthy compared to more trustworthy-looking faces, effects of appearance-based trustworthiness were likely too small and not consistent enough to yield a significant difference in CBP tests. Notably, effects of facial trustworthiness in the literature are generally rather heterogeneous (see Introduction section 1.2) and likely depend strongly on the specific set of stimuli used.

Overall, trustworthiness and emotional expression evaluations indicate successful manipulations of both factors, with independent contributions of affective knowledge and facial trustworthiness. In ERPs, a typical EPN effect was observed for affective knowledge, with enhanced negative mean amplitudes for faces associated with negative as compared to neutral knowledge, while no systematic differences in brain activity corresponding to facial trustworthiness were found in the examined components.

### 4.3. Effects of Affective Knowledge on Visual Awareness: Behavioral Results

In the attentional blink task, visual awareness under conditions of reduced attention was enhanced for faces associated with negative as compared to neutral knowledge. Since the assignment of faces to affective knowledge condition was counterbalanced across participants, low-level visual differences cannot explain this finding. Replicating a previous report (Eiserbeck & Abdel Rahman, 2020), the effect depended on the subjective visibility rating and was only observed for the strict hit criterion (hit = correct gender classification and at least visibility rating 3, “strong impression”; miss = all other trials), not for the liberal criterion of at least a slight impression (hit = correct gender classification and at least visibility rating 2, “slight impression”; miss = all other trials). This result pattern indicates that the intensity or quality of the percept—rather than only the precision with which the objective (gender classification) task is solved—is influenced by affective knowledge (see Eiserbeck & Abdel Rahman, 2020; Fazekas & Overgaard, 2018). This is in line with findings indicating gradual variations of stimulus awareness during the attentional blink in general (Cohen et al., 2023; Eiserbeck et al., 2022; Nieuwenhuis & de Kleijn, 2011; Roth-Paysen et al., 2022) and specifically in regard to emotional manipulations (Keefe & Zald, 2022).

### 4.4. Effects of Affective Knowledge on Visual Awareness: ERPs

The behavioral effect was accompanied by an early ERP modulation in the EPN time range, with a corresponding cluster from 210 to 240 ms after T2 onset at posterior electrodes. The mean amplitude in this time range and region of interest had a higher predictive value for face awareness in the negative compared to the neutral knowledge condition. This difference was present for both liberal and strict hit criterion, unlike in the behavior data, where an effect was only found for the strict hit criterion. This discrepancy between behavioral and ERP data might be due to the higher sensitivity and more comprehensive mapping of differences in the neurophysiological measure: ERPs enable a tracking of the continuous unfolding of brain processing over time, whereas behavioral measures represent a combined result of different neural processes. Similar discrepancies, with attentional effects being observable in ERPs and not in behavioral data, have been reported in previous studies (Kappenman et al., 2013, 2014; Xu et al., 2016).

The results point to slightly earlier differences between affective knowledge conditions (in interaction with awareness) in the attentional blink task compared to the evaluations. This pattern of results could be explained by the different task demands as well as differences in the contrast that is considered in the analyses: During the evaluations, the images were shown comparatively long (1 s) and one by one. In this task, attentional resources were sufficiently available to enable full conscious processing of all stimuli, i.e., no early selection based on emotional relevance was necessary. In the attentional blink task, the images were presented only very briefly (117 ms), preceded and followed by distractor stimuli, leading to the intended restrictions in conscious awareness. Here, the influence of affective knowledge was observable at an earlier temporal stage presumably since it was relevant already in determining the significance of a stimulus for conscious perception in the first place.

To put the observed knowledge effect in context, the overall temporal course of processing during the attentional blink should be considered. To this end, we also analyzed overall differences between hit and miss trials during the VAN and LP time range. Significant differences were found, with corresponding clusters spanning the whole tested time ranges (150-350 ms for VAN, 400-800 ms for LP). In line with this, a companion article (Eiserbeck et al., 2022) based on the same attentional blink data addressed the overall time course of the access to visual awareness in more detail and found graded differences based on reported visibility starting at around 150 ms in the N1 component and continuing in the N2 and P3 time range. As described earlier in regard to the interpretation of CBP test results, the onset of the cluster corresponding to the social-emotional knowledge effect in the present study (210 ms) cannot be interpreted as the actual onset of the effect, which may already occur earlier in time. However, it can be concluded that the effect is found soon after the start of overall differences in brain activity correlating with stimulus awareness. Specifically, the social-emotional effect emerges during the time range of the broader VAN component, which has been characterized as the most consistent ERP correlate of visual consciousness across different paradigms and is assumed to reflect phenomenal consciousness, i.e., the subjective experience of seeing (Förster et al., 2020; Koivisto & Revonsuo, 2010). The occurrence of the emotional effect within this processing phase aligns with the notion that the resulting perceptual experience is modulated based on the social-emotional relevance of a face acquired through verbal information. Thus, awareness of the target stimulus is characterized by broad activity differences (the whole VAN time range) that may result from trial-by-trial differences in attentional engagement to T2. Activity during this time range might reflect the buildup of conscious content from a coarse to a more detailed percept (see Aru & Bachmann, 2017; Campana et al., 2016). Relatively early in this process, the emotional knowledge associated with a stimulus modulates this buildup of conscious content.

Follow-up analyses investigating the direction of effects, revealed awareness (hit-miss) differences during the observed cluster in the negative, but not the neutral knowledge condition. In line with this, in GLMM analyses, which enabled the prediction of awareness (hit/miss) based on all relevant variables and their interaction within one model, an increased predictive value of the mean ROI amplitude during the 210-240 ms time was found for the negative compared to the neutral knowledge condition. This pattern of results may be attributed to heightened top-down processing: Following or concurrent with the processing of facial identity within the N170 time range (Heisz et al., 2006; Hinojosa et al., 2015), the associated emotional value is rapidly accessed and exerts an influence on the quality of the resulting conscious percept. The activity demonstrates systematic effects on awareness specifically for faces associated with negative information. In contrast, for faces with neutral information, the activity during this region of interest and time range exhibits less systematic prediction value, in line with the lower diagnostic relevance of neutral information concerning potential threats. Instead, the processing of neutral faces may be more strongly influenced by general effects of trial-by-trial differences in attention.

When comparing the activity following T2-onset solely in *hit* trials for faces associated with neutral and negative knowledge, no significant differences were observed. This indicates that the observed effect cannot be explained merely by knowledge effects for faces that have already been consciously perceived (i.e., where the transition to conscious perception has already been completed). Combined with the observed enhanced awareness of faces associated with negative knowledge in the behavioral data, this suggests an intertwining of social-affective knowledge with visual awareness—rather than effects of knowledge depending on conscious perception. In other words: Social-affective relevance seems to play an active role in shaping what is ultimately perceived.

Unexpectedly, in the comparison between neutral and negative knowledge in *miss* trials, significant differences were observed, with more positive amplitudes for negative as compared to neutral knowledge miss trials. The direction of this effect goes in the opposite direction of our general expectation for the EPN component, for which we hypothesized an enhanced negativity in the negative as compared to the neutral knowledge condition. However, for the specific case of miss trials, we did not have any expectations in regard to knowledge differences, and in general the observed pattern does not contradict any predictions (as it was not accompanied by a significant main effect of knowledge in the same direction). The effect may be indicative of opposing directions within the negative knowledge condition. While the behavioral effect indicated an overall awareness advantage for negative faces, on the level of single items negative knowledge may have differential effects. That is, similar to different coping styles in the presence of threat (e.g., “fight or flight”), depending on specifics of the associated negative information in combination with the respective face and characteristics of the participants (e.g., individual previous experiences), some negative knowledge items (the majority) may lead to attentional enhancement, while others may lead to an avoidance/disregard of the visual information. Thus, negative knowledge may lead to stronger reactions in both directions compared to neutral knowledge. The enhanced positivity in negative miss trials compared to neutral miss trials may thus be reflective of a more effective disregard of specific items. Future research could further investigate this direction by focusing on participant- and item-specific differences in the response to negative social knowledge.

While overall awareness differences also showed strong effects during the LP time range (also see Eiserbeck et al., 2022), no emotion-specific effects were found during this stage. This may be explained by the task relevance of specific aspects: In the attentional blink task, face detection itself was task-relevant for the male/female categorization task. This may explain the presence of overall awareness effects during the LP time range in contrast to a recent attentional blink study where the task relevance of stimuli was unknown during stimulus presentation (Dellert et al., 2022). The emotional value of stimuli, on the other hand, was not task-relevant in our attentional blink task, which may explain the lack of LPP/LP differences based on emotional knowledge (for an examination of the role of task relevance on LPP effects, see Schindler & Straube, 2020). To directly test this, future studies could compare the impact of different task requirements on the LP during the attentional blink by including an alternative task requiring an emotional judgment.

To conclude, the findings indicate that social-affective knowledge associated with a person affects perception-related processing during a phase of attentional enhancement for conscious awareness. As a result, faces associated with negative compared to neutral information are prioritized for visual awareness. These findings replicate and extend results from a previous behavioral study (Eiserbeck & Abdel Rahman, 2020). To the best of our knowledge, these studies are the first to report influences of social-affective knowledge on the visual consciousness of faces, whereas (apart from the discussed initial evidence from E. Anderson et al., 2011) this could not be shown in previous studies with other paradigms (binocular rivalry and breaking continuous flash suppression; Rabovsky et al., 2016; Stein et al., 2017). These differing results may be due to differences in the suppression techniques utilized in the paradigms (for a comparison of underlying mechanisms, see, e.g., Kanai et al., 2010): Binocular rivalry and continuous flash suppression rely on interocular suppression—a suppression of low-level sensory signals—and it is not yet clear whether or to what extent higher-level (e.g., emotional) processing of stimuli is possible under these conditions (Moors et al., 2017, 2019; Sklar et al., 2018). The attentional blink, on the other hand, is characterized by attentional blindness (i.e., low-level signals cannot be accessed despite being available) and may enable processing up to a conceptual level (Martens & Wyble, 2010).

### 4.5. Facial Trustworthiness and Visual Awareness

In contrast to the affective knowledge effects, we found no clear evidence for an impact of appearance-based trustworthiness inferences on visual consciousness in the attentional blink. In the behavioral data, only a non-significant trend for a main effect of appearance was observed (whereas in one model additionally including the ERP amplitude in the EPN time range, the p-value just exceeded the threshold for statistical significance, see Appendix Table A3.1). This trend followed the direction postulated in the hypotheses: Less trustworthy faces showed a tendency for enhanced awareness under conditions of reduced attention. The interaction between appearance and knowledge did not reach statistical significance. Interestingly, investigating appearance effects separately within each knowledge condition indicated an influence of appearance on awareness in the neutral but not in the negative knowledge condition (strict hit criterion; see Appendix Table A3.2), possibly pointing towards a stronger influence of appearance-based trustworthiness when associated knowledge is less diagnostic (neutral knowledge condition). This could be an interesting aspect to further test in future work. However, as the overall interaction did not reach significance, no definite conclusions can be drawn from this pattern of results.

The ERP analyses revealed a significant main effect of appearance in the EPN time range, with more positive amplitude values for untrustworthy as compared to trustworthy looking faces. The corresponding cluster was observed between 180 to 206 ms. The effect showed the opposite direction of what would have been expected for a main effect of appearance, namely, an enhanced negativity for more trustworthy faces. While the effect reflects small differences based on appearance during the attentional blink task, no interaction of appearance and awareness was found, indicating that the processing was not relevant in regard to the access to awareness. Furthermore, as a main effect of appearance was found during the attentional blink task, but not during viewing of the faces in the rating task, the observed effect may be indicative of mere visual differences (which can be expected to have a stronger impact in a task focused on detection) rather than differences in attributed affective value. We conclude that, even though facial trustworthiness affects explicit and conscious evaluations of persons and expressions and may have been processed to a certain degree in the attentional blink task, its influence is very limited, in line with a previous report comparing knowledge- and appearance-based trustworthiness effects on visual consciousness (Eiserbeck & Abdel Rahman, 2020). Electrophysiological results yielded no evidence for connections between facial trustworthiness and awareness.

### 4.6. Limitations and Future Directions

As already described, the CBP analyses used in the present study come with advantages and with disadvantages: While they allowed the examination of effects within the EPN and LPP time range and regions of interest with flexibility in regard to the exact latency and topography of the effects, small effects may be hard to uncover. Furthermore, CBP analyses in the present implementation are conducted based on condition averages per participant, with differing numbers of trials per condition entering the attentional blink analyses based on participants’ task performance. We confirmed and further examined the results by running additional single-trial analyses with linear mixed models based on mean amplitudes across the observed cluster, which enabled us to also take into account random effect variances for participants and items. A combination of both methods (running linear mixed models for every time point × electrode combination, and using CBP tests on the obtained coefficients) was not technically feasible as yet, but with advances in methods (e.g., Visalli et al., 2023) this might be an option in future studies in order to combine the advantages of both methods.

The used emotional knowledge manipulation—the presentation and learning of one-sentence affective information for each face—has been used in a number of previous studies, which observed effects of affective information (Luo et al., 2016; Morel et al., 2012; Verosky et al., 2018; Verosky & Todorov, 2013; Xu et al., 2016). Manipulation checks revealed a successful manipulation of affective knowledge in the present study, yielding the expected behavioral results and EPN differences during conscious perception of faces in the rating task. Yet, EPN differences during the rating task as well as awareness differences during the attentional blink based on affective knowledge were rather small. Future experiments could aim to induce stronger knowledge manipulations which may also enhance effects on awareness, for example by enhancing the ecological validity through an embedding of faces and affective information in the context of newspaper articles during the learning phase (Baum & Abdel Rahman, 2021b, 2021a; Schindler et al., 2021).

### 4.7. Conclusions

The present study provides evidence for an influence of social-affective knowledge on the visual awareness of faces. Replicating previous findings (Eiserbeck & Abdel Rahman, 2020), faces associated with negative information had a higher chance of being selected for enhanced conscious processing than faces associated with relatively neutral information. Cluster-based permutation tests revealed a connection between perception- and attention-related processing during the time range of the EPN (latency of the observed cluster: 210-240 ms). Specifically, we observed a higher predictive value of posterior ERP amplitudes in determining awareness of faces associated with negative compared to neutral information. Our findings suggest that social-affective knowledge about individuals can influence to what degree visual facial information becomes available for conscious processing, providing an important basis for social perception. Beyond descriptive trends in the behavioral data, no evidence for an effect of facial trustworthiness on visual awareness was observed.

## Data and code availability

Data and analysis scripts will be made publicly available upon peer-reviewed publication.

## Acknowledgments

We thank Guido Kiecker for technical support and Friedrich Eiserbeck, Franziska Glogau, Maja Heitmann, Hannah Kaube, Kirsten Stark, and Nura Völk for their help with data collection.

## Appendix A1: Stimulus Material

**Table A1.** Sentences Containing Social-Affective Information Used in the Study (English Translations of Used German Sentences)

| Knowledge category | Sentence | Valence rating (1-7) | SD | Arousal rating (1-7) | SD |
| --- | --- | --- | --- | --- | --- |
| Negative | He made a lot of money with child pornography. | 1.10 | 0.31 | 5.50 | 1.73 |
| Negative | This woman set fire to a refugee shelter. | 1.20 | 0.41 | 5.75 | 1.45 |
| Negative | He fired into a crowd near a church. | 1.35 | 0.67 | 5.75 | 1.59 |
| Negative | This man has killed baby monkeys in a zoo. | 1.37 | 0.60 | 5.25 | 1.62 |
| Negative | This man beat his wife regularly. | 1.70 | 1.08 | 5.20 | 1.40 |
| Negative | This woman cheated her best friend out of a lot of money. | 1.70 | 1.08 | 4.60 | 1.82 |
| Negative | He threatened a shop assistant with a knife. | 1.75 | 0.79 | 4.70 | 1.53 |
| Negative | She sabotaged the brake lines on her neighbors' bikes. | 1.80 | 0.70 | 4.60 | 1.64 |
| Negative | She ran over a man and committed a hit-and-run. | 1.85 | 0.99 | 4.85 | 1.63 |
| Negative | She made fun of a child with a disability. | 1.85 | 0.81 | 4.55 | 1.73 |
| Negative | This man slapped his colleague for no reason. | 2.20 | 0.83 | 4.35 | 1.84 |
| Negative | This woman pretends to be a cleaning lady to steal from her customers. | 2.21 | 0.71 | 4.15 | 1.50 |
| Neutral | She took the change from the shop assistant. | 3.85 | 0.88 | 1.45 | 1.19 |
| Neutral | She described the haircut to the hairdresser. | 3.95 | 0.60 | 1.35 | 0.49 |
| Neutral | He showed a technician the connection. | 3.95 | 0.76 | 1.30 | 0.57 |
| Neutral | She handed in forms to the clerk. | 4.00 | 0.46 | 1.50 | 1.19 |
| Neutral | This man made an appointment with an eye doctor. | 4.00 | 0.79 | 1.20 | 0.41 |
| Neutral | This man asked a waiter for the menu. | 4.00 | 0.56 | 1.15 | 0.37 |
| Neutral | This man spoke to his employer. | 4.05 | 0.22 | 1.60 | 1.39 |
| Neutral | This woman took the elevator with a neighbor. | 4.05 | 0.76 | 1.40 | 0.75 |
| Neutral | He was watching an old western on night television. | 4.10 | 0.45 | 1.45 | 1.00 |
| Neutral | He received a registered letter from the mailman. | 4.10 | 0.72 | 1.20 | 0.62 |
| Neutral | This woman left her dress at the tailor's. | 4.15 | 0.49 | 1.40 | 0.99 |
| Neutral | This woman opened the door for a salesman. | 4.25 | 0.85 | 1.40 | 0.60 |
| Positive (Filler) | This man has taken stranded seals back into the sea. | 6.20 | 0.95 | 4.65 | 1.50 |
| Positive (Filler) | This woman is letting a refugee stay with her for free. | 6.30 | 1.22 | 4.35 | 1.66 |
| Positive (Filler) | This woman is protecting endangered species in the jungle from wild predators. | 6.30 | 0.73 | 4.10 | 1.80 |
| Positive (Filler) | This man carried an injured climber down into the valley. | 6.40 | 0.75 | 5.05 | 1.50 |
| Positive (Filler) | This woman donated a kidney to her sick sister. | 6.53 | 0.77 | 4.65 | 1.50 |
| Positive (Filler) | This man rescued an injured woman from her crashed car. | 6.60 | 0.68 | 4.70 | 1.59 |

## Appendix A2: Differences to the Pre-Registrations

The hypotheses and methods of this study were pre-registered using the Open Science Framework (OSF) and can be accessed under https://osf.io/us754 (pre-registration 1; for the aspect of affective knowledge) and https://osf.io/2yspe (pre-registration 2; for the aspect of facial trustworthiness and its interaction with affective knowledge).

### A2.1 ERP Analyses in Post-Learning Rating Phase

To avoid the issue of double-dipping (identifying an effect based on the data and then testing it), we used cluster-based permutation tests for the analyses of EPN and LPP effects during the post-learning rating phase rather than linear mixed model analyses as stated in the pre-registration.

### A2.2 ERP Analyses in Attentional Blink Task

#### A2.2.1 Methods and Considerations

Since ERPs in the attentional blink task are noisier due to the rapid serial visual presentation of multiple images, we had planned to use the rating task as a localizer task. The idea was to describe EPN effects during the rating task and then analyze the same regions of interest and time range in the attentional blink task to examine the presence of EPN effects after T2-face presentation during short lag attentional blink trials. To this end, based on the topographical and temporal distribution of the cluster corresponding to a significant knowledge effect in the rating task, single-trial mean amplitudes were obtained and analyzed as the dependent variable in a linear mixed model containing the factors knowledge (neutral/negative) and appearance (continuous predictor), with effect coding applied for the factor knowledge (neutral: -0.5, negative: 0.5) and mean-centering the predictor appearance. This approach was unsuccessful, likely due to task differences and interactions with awareness. We report the results of this analysis in the following paragraph. Additional analyses were conducted using cluster-based permutation tests, as described in the article.

#### A2.2.2 Results for Planned Analyses

In correspondence to the distribution of the main effect of affective knowledge observed in the rating task, the LMM analysis was focused on the time range of 270 to 288 ms and electrodes P4, P6, P8, PO3, POz, PO4, PO8, PO10, Oz and O2. No significant effects in the prediction of mean amplitudes were observed for either affective knowledge (*b* = 0.15, *t*[45.26] = 1.04, *p* = .302), appearance (*b* = -0.13, *t*[22.81] = -1.07, *p* = .297) or an interaction between affective knowledge and appearance (*b* = -0.02, *t*[3172.42] = -0.10, *p* = .923).

### A2.3 Exploratory Analyses

Pre-registration 2 also lists the P1 component to investigate in exploratory analyses (specifically for facial trustworthiness). We decided not to focus on this component in order to not exceed the scope of the present manuscript, and because such early modulations are more likely to reflect visual differences rather than differences in emotional relevance, which are of interest in the present manuscript.

## Appendix A3: Linear Mixed Model Output

**Table A3.1.**
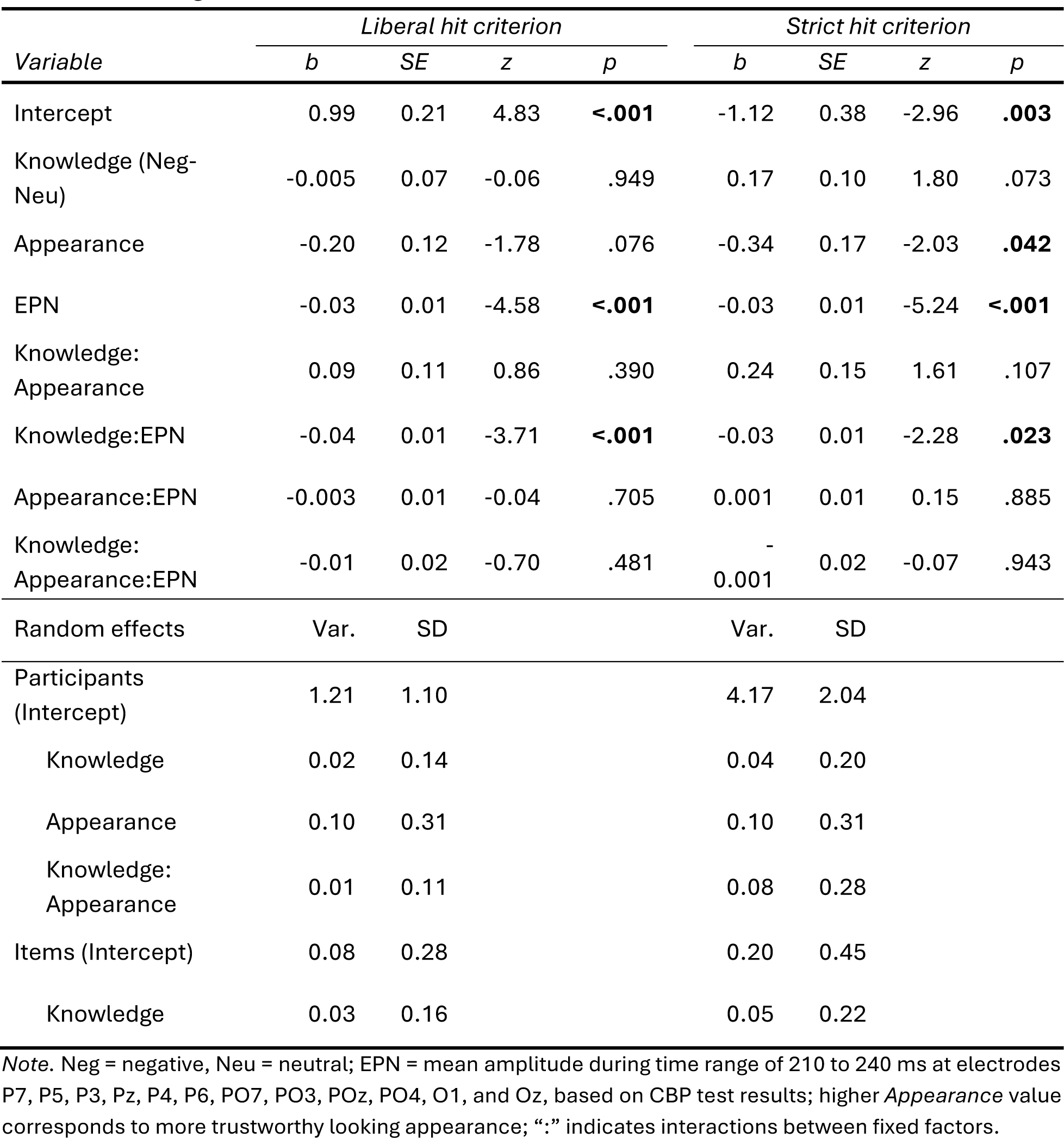
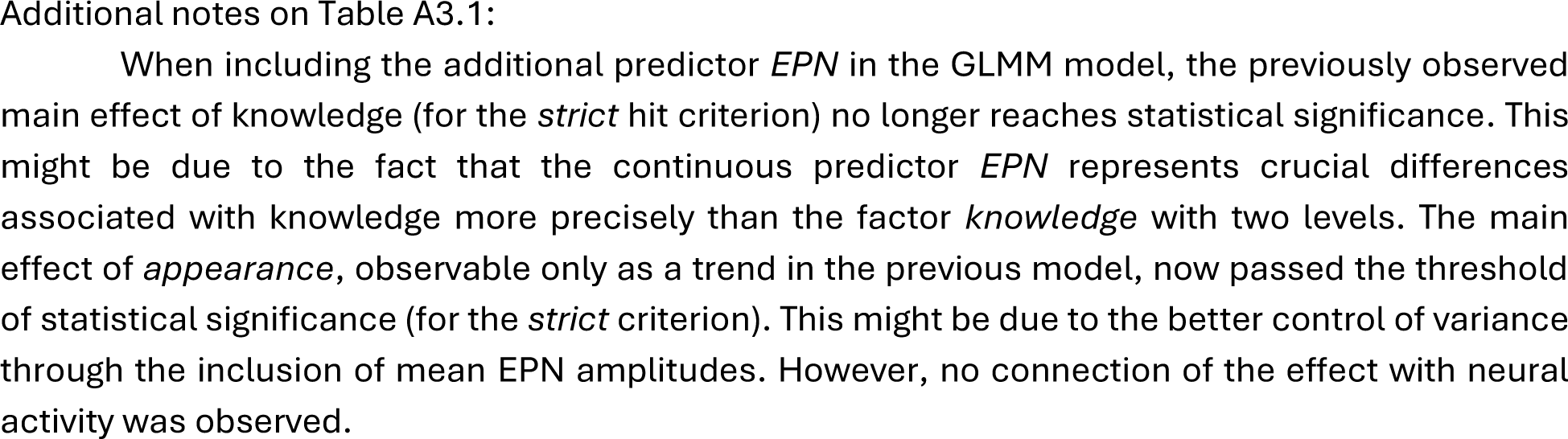
GLMM Statistics for Analysis of Short Lag Attentional Blink Trials Including Mean Amplitude During the EPN Time Range as a Predictor.

**Table A3.2.** GLMM Statistics for Analysis of Short Lag Attentional Blink Trials Including Mean Amplitude During the EPN Time Range as a Predictor (Appearance and Mean Amplitude Nested Within Knowledge Condition).

| Variable | Liberal hit criterion |  |  |  | Strict hit criterion |  |  |  |
| --- | --- | --- | --- | --- | --- | --- | --- | --- |
|  | *b* | SE | *z* | *p* | *b* | SE | *z* | *p* |
| Intercept | 0.99 | 0.21 | 4.83 | **<.001** | -1.12 | 0.38 | -2.96 | **.003** |
| Knowledge<br>(Neg-Neu) | -0.005 | 0.07 | -0.06 | .949 | 0.17 | 0.10 | 1.80 | .073 |
| Neu/EPN | -0.01 | 0.01 | -0.71 | .476 | -0.02 | 0.01 | -2.17 | **.030** |
| Neg/EPN | -0.05 | 0.01 | -5.92 | **<.001** | -0.05 | 0.01 | -5.43 | **<.001** |
| Neu/Appearance | -0.25 | 0.13 | -1.92 | .055 | -0.46 | 0.16 | -2.80 | **.005** |
| Neg/Appearance | -0.16 | 0.12 | -1.30 | .205 | -0.22 | 0.20 | -1.10 | .271 |
| Neu/(EPN:<br>Appearance) | 0.002 | 0.01 | 0.23 | .820 | 0.002 | 0.01 | 0.15 | .882 |
| Neg/(EPN:<br>Appearance) | -0.01 | 0.01 | -0.75 | .453 | 0.001 | 0.01 | 0.06 | .956 |
| Random effects | Var. | SD |  |  | Var. | SD |  |  |
| Participants<br>(Intercept) | 1.21 | 1.10 |  |  | 4.17 | 2.04 |  |  |
| Knowledge | 0.02 | 0.14 |  |  | 0.04 | 0.20 |  |  |
| Appearance | 0.10 | 0.31 |  |  | 0.10 | 0.31 |  |  |
| Knowledge:<br>Appearance | 0.01 | 0.11 |  |  | 0.08 | 0.28 |  |  |
| Items<br>(Intercept) | 0.08 | 0.28 |  |  | 0.20 | 0.45 |  |  |
| Knowledge | 0.03 | 0.16 |  |  | 0.05 | 0.22 |  |  |

*Note.* Neg = negative, Neu = neutral; EPN = mean amplitude during time range of 210 to 240 ms at electrodes P7, P5, P3, Pz, P4, P6, PO7, PO3, POz, PO4, O1, and Oz, based on CBP test results; higher *Appearance* value corresponds to more trustworthy looking appearance; “:” indicates interactions between fixed factors, “/” indicates nesting of fixed factors.

## Appendix A4: Control Analyses – Attentional Blink Task

In control analyses, participants with less than 10 ERP trials in any of the individual knowledge×awareness or appearance×awareness conditions in short lag attentional blink trials were excluded (N = 14), leaving 18 data sets for analyses.

### A4.1. Behavioral Results

**Table A4.1.1.** Control Analyses (N = 18): GLMM Statistics for Analysis of Short Lag Trials.

| Variable | Liberal hit criterion |  |  |  | Strict hit criterion |  |  |  |
| --- | --- | --- | --- | --- | --- | --- | --- | --- |
|  | *b* | *SE* | *z* | *p* | *b* | *SE* | *z* | *p* |
| Intercept | 0.92 | 0.22 | 4.27 | **<.001** | -0.81 | 0.28 | -2.88 | **.004** |
| Knowledge (Neg-Neu) | -0.006 | 0.09 | -0.07 | .946 | 0.21 | 0.08 | 2.53 | **.012** |
| Appearance | -0.19 | 0.14 | -1.46 | .146 | -0.35 | 0.18 | -1.94 | .052 |
| Knowledge:Appearance | 0.19 | 0.14 | 1.31 | .189 | 0.22 | 0.13 | 1.73 | .084 |
| Random effects | Var. | SD |  |  | Var. | SD |  |  |
| Participants (Intercept) | 0.73 | 0.85 |  |  | 1.22 | 1.10 |  |  |
| Knowledge | 0.001 | 0.04 |  |  | 0.02 | 0.13 |  |  |
| Appearance | 0.07 | 0.27 |  |  | 0.05 | 0.22 |  |  |
| Knowledge:Appearance | 0.01 | 0.10 |  |  | 0.03 | 0.17 |  |  |
| Items (Intercept) | 0.11 | 0.33 |  |  | 0.26 | 0.51 |  |  |
| Knowledge | 0.06 | 0.25 |  |  | 0.02 | 0.14 |  |  |

*Note.* Neg = negative, Neu = neutral; higher *Appearance* value corresponds to a more trustworthy looking appearance; “:” indicates interactions between fixed factors.

### A4.2. ERP results

In the EPN time range, CBP tests revealed a significant interaction of awareness and affective knowledge for the liberal criterion (*p* = .010), indicating an enhanced negative difference between hit and miss trials in the negative as compared to the neutral knowledge condition (see Figure A4.2). The corresponding cluster was observed from 190 to 228 ms at electrodes P7, P5, P3, Pz, P4, P6, PO9, PO7, PO3, POz, PO4, PO8, O1, Oz, and O2. For the strict hit criterion, the interaction of awareness and affective knowledge failed to reach significance (smallest cluster-level *p* = .064). No interaction between appearance and awareness was found (cluster-level *p* = .375).

For comparability and due to the similar cluster distribution, in additional linear mixed model analyses the same time range and electrodes as in the main analyses were used: 210 to 240 ms at electrodes P7, P5, P3, Pz, P4, P6, PO7, PO3, POz, PO4, O1, and Oz. The model specifications were the same as in the main analyses. The models revealed a significant interaction between affective knowledge and mean amplitude for the liberal hit criterion (*b* = -0.04, *z* = -2.62, *p* = .009). Nested models showed that the mean amplitude only has predictive value for determining a trial as a hit or miss in the negative knowledge condition (*b* = -0.04, *z* = -4.38, *p* < .001) and not in the neutral knowledge condition (*b* = -0.01, *z* = -0.64, *p* = .525). For the strict hit criterion, the interaction did not reach significance (*b* = -0.02, *z* = -1.67, *p* = .095), but descriptively the pattern observed in the corresponding nested model matched the pattern in the main analyses, where a larger effect of the mean amplitude in the negative condition had been found. For full model output, see Appendix A4.2.1 and A4.2.2.

**Table A4.2.1.**
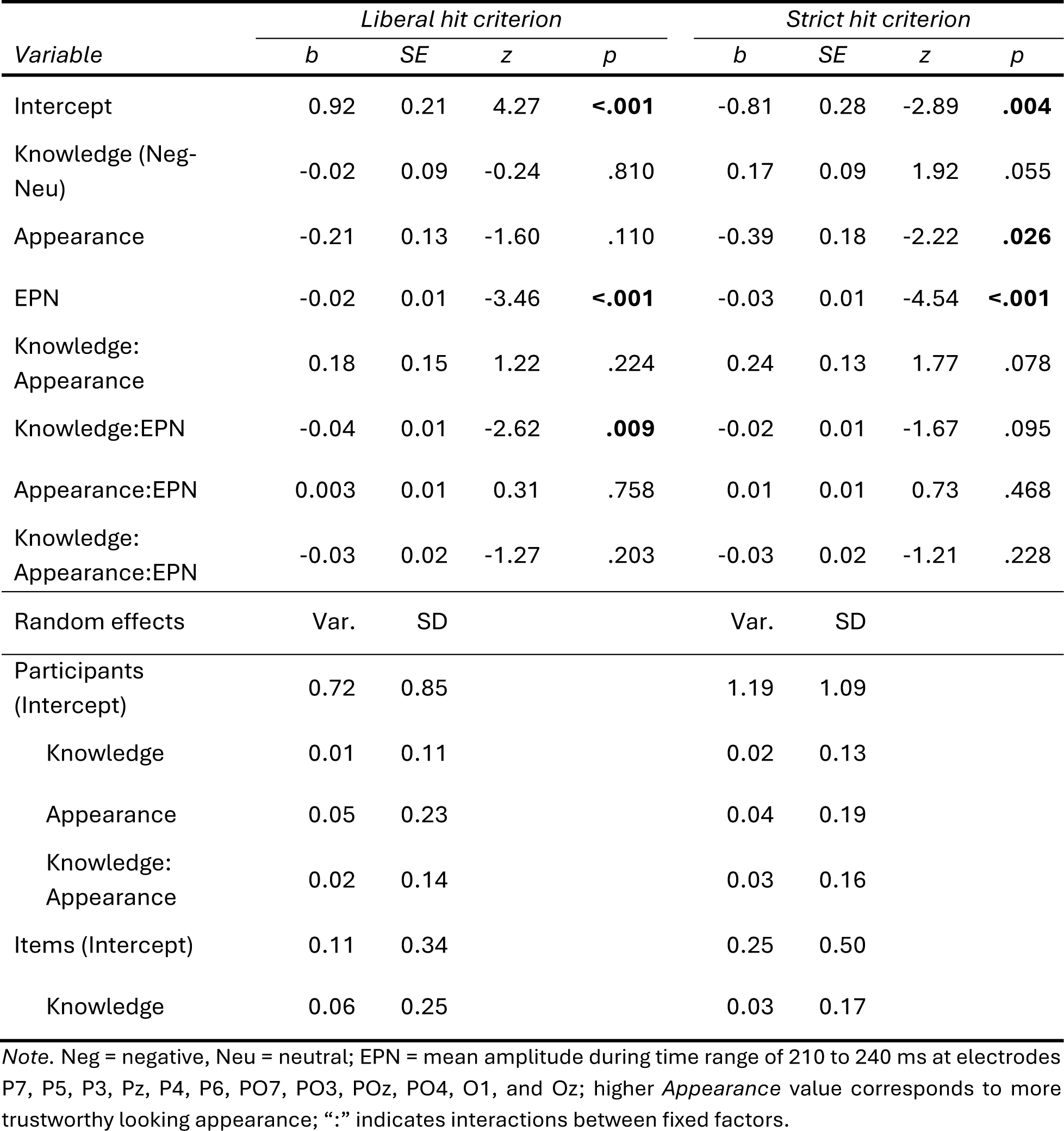
Control Analyses (N = 18): GLMM Statistics for Analysis of Short Lag Attentional Blink Trials Including Mean Amplitude During the EPN Time Range as a Predictor.

| *Variable* | *Liberal hit criterion* |  |  |  | *Strict hit criterion* |  |  |  |
| --- | --- | --- | --- | --- | --- | --- | --- | --- |
|  | *b* | *SE* | *z* | *p* | *b* | *SE* | *z* | *p* |
| Intercept | 0.92 | 0.21 | 4.27 | **<.001** | -0.81 | 0.28 | -2.89 | **.004** |
| Knowledge (Neg-Neu) | -0.02 | 0.09 | -0.24 | .810 | 0.17 | 0.09 | 1.92 | .055 |
| Appearance | -0.21 | 0.13 | -1.60 | .110 | -0.39 | 0.18 | -2.22 | **.026** |
| EPN | -0.02 | 0.01 | -3.46 | **<.001** | -0.03 | 0.01 | -4.54 | **<.001** |
| Knowledge: Appearance | 0.18 | 0.15 | 1.22 | .224 | 0.24 | 0.13 | 1.77 | .078 |
| Knowledge:EPN | -0.04 | 0.01 | -2.62 | **.009** | -0.02 | 0.01 | -1.67 | .095 |
| Appearance:EPN | 0.003 | 0.01 | 0.31 | .758 | 0.01 | 0.01 | 0.73 | .468 |
| Knowledge: Appearance:EPN | -0.03 | 0.02 | -1.27 | .203 | -0.03 | 0.02 | -1.21 | .228 |
| Random effects | Var. | SD |  |  | Var. | SD |  |  |
| Participants (Intercept) | 0.72 | 0.85 |  |  | 1.19 | 1.09 |  |  |
| Knowledge | 0.01 | 0.11 |  |  | 0.02 | 0.13 |  |  |
| Appearance | 0.05 | 0.23 |  |  | 0.04 | 0.19 |  |  |
| Knowledge: Appearance | 0.02 | 0.14 |  |  | 0.03 | 0.16 |  |  |
| Items (Intercept) | 0.11 | 0.34 |  |  | 0.25 | 0.50 |  |  |
| Knowledge | 0.06 | 0.25 |  |  | 0.03 | 0.17 |  |  |

*Note.* Neg = negative, Neu = neutral; EPN = mean amplitude during time range of 210 to 240 ms at electrodes P7, P5, P3, Pz, P4, P6, PO7, PO3, POz, PO4, O1, and Oz; higher *Appearance* value corresponds to more trustworthy looking appearance; “:” indicates interactions between fixed factors.

**Table A4.2.2.** Control Analyses (N = 18): GLMM Statistics for Analysis of Short Lag Attentional Blink Trials Including Mean Amplitude During the EPN Time Range as a Predictor (Appearance and Mean Amplitude Nested Within Knowledge Condition).

| Variable | Liberal hit criterion |  |  |  | Strict hit criterion |  |  |  |
| --- | --- | --- | --- | --- | --- | --- | --- | --- |
|  | *b* | SE | *z* | *p* | *b* | SE | *z* | *p* |
| Intercept | 0.92 | 0.21 | 4.27 | **<.001** | -0.81 | 0.28 | -2.89 | **.004** |
| Knowledge (Neg-Neu) | -0.02 | 0.09 | -0.24 | .811 | 0.17 | 0.09 | 1.92 | .055 |
| Neu/EPN | -0.01 | 0.01 | -0.64 | .525 | -0.02 | 0.01 | -2.03 | **.043** |
| Neg/EPN | -0.04 | 0.01 | -4.38 | **<.001** | -0.04 | 0.01 | -4.56 | **<.001** |
| Neu/Appearance | -0.30 | 0.16 | -1.91 | .057 | -0.51 | 0.17 | -2.95 | **.003** |
| Neg/Appearance | -0.12 | 0.15 | -0.85 | .397 | -0.27 | 0.20 | -1.35 | .177 |
| Neu/(EPN: Appearance) | 0.02 | 0.02 | 1.11 | .269 | 0.02 | 0.02 | 1.33 | .185 |
| Neg/(EPN: Appearance) | -0.01 | 0.02 | -0.68 | .495 | -0.01 | 0.01 | -0.34 | .734 |
| Random effects | Var. | SD |  |  | Var. | SD |  |  |
| Participants (Intercept) | 0.72 | 0.85 |  |  | 1.19 | 1.09 |  |  |
| Knowledge | 0.01 | 0.11 |  |  | 0.02 | 0.13 |  |  |
| Appearance | 0.05 | 0.23 |  |  | 0.04 | 0.19 |  |  |
| Knowledge: Appearance | 0.02 | 0.14 |  |  | 0.03 | 0.16 |  |  |
| Items (Intercept) | 0.11 | 0.34 |  |  | 0.25 | 0.50 |  |  |
| Knowledge | 0.06 | 0.25 |  |  | 0.03 | 0.17 |  |  |

*Note.* Neg = negative, Neu = neutral; EPN = mean amplitude during time range of 210 to 240 ms at electrodes P7, P5, P3, Pz, P4, P6, PO7, PO3, POz, PO4, O1, and Oz; higher *Appearance* value corresponds to more trustworthy looking appearance; “:” indicates interactions between fixed factors, “/” indicates nesting of fixed factors.

**Figure A4.1.**
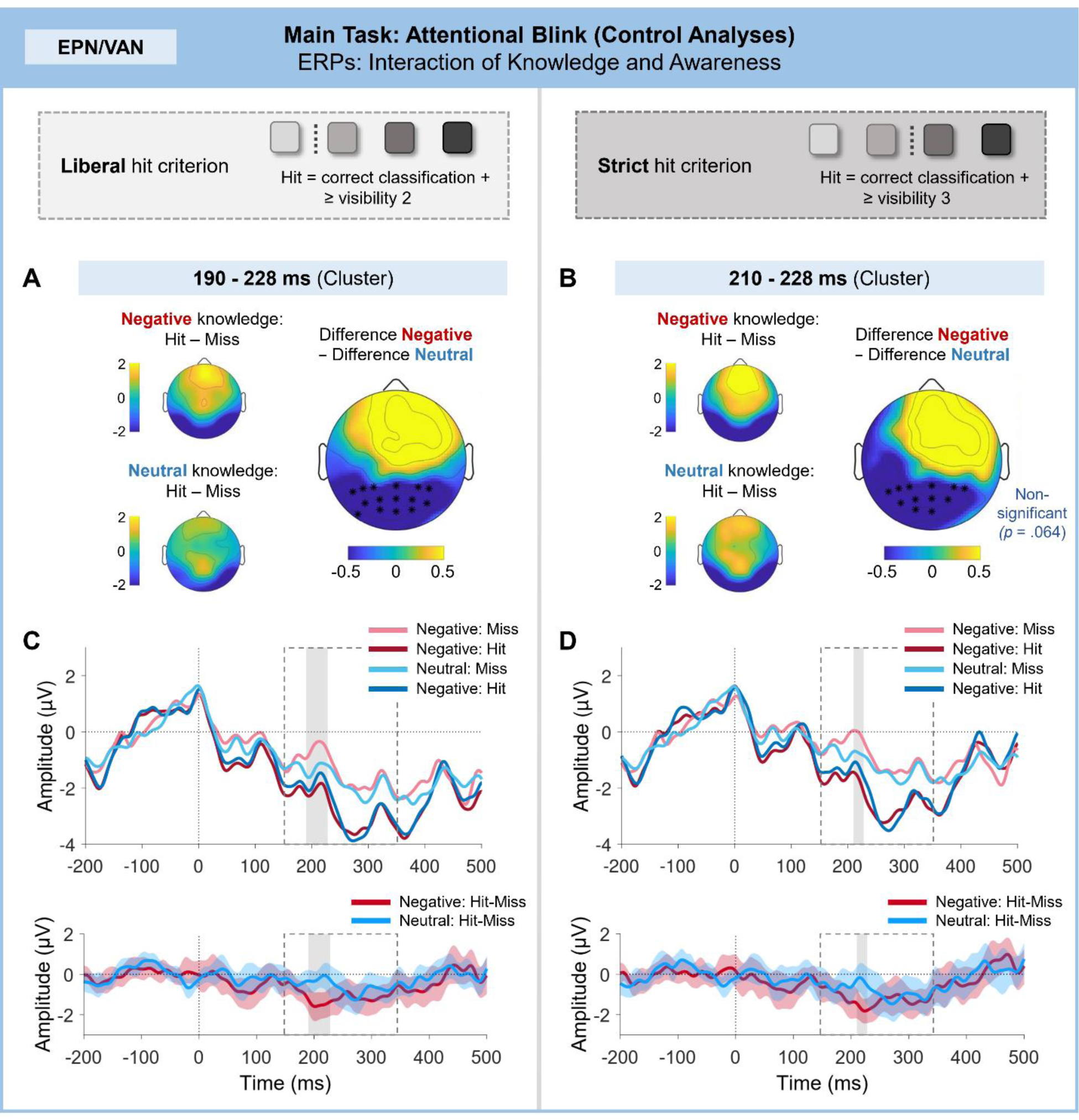
ERP differences during the EPN/VAN time range in the attentional blink for the control analyses: Interaction of knowledge and awareness. Cluster-based permutation tests revealed a similar pattern of results as in the main analyses. A significant interaction of knowledge and awareness in the EPN time range was found for the liberal hit criterion. For the strict hit criterion, the interaction effect failed to reach statistical significance (*p* = .064) but descriptively the pattern matched that of the main analyses. The corresponding (non-significant) cluster is displayed for comparison. (A), (B) The smaller topographies on the left show the difference between Hit and Miss trials in the negative and neutral knowledge conditions in the interval of the cluster. The topography on the right shows the corresponding double difference (awareness difference in negative condition – awareness difference in neutral condition). Asterisks mark the electrodes included in the cluster. (C), (D) Grand-average event-related potential for each knowledge × awareness condition, based on pooled activity across electrode sites included in the cluster (upper panel), and corresponding difference curves with 95% bootstrap confidence intervals for Hit-Miss trials in the negative and neutral knowledge conditions (lower panel). The dashed frame indicates the time range included in the CBP test; the gray frame indicates the temporal extent of the cluster.

## Appendix A5: Additional Figures

**Figure A5.1.**
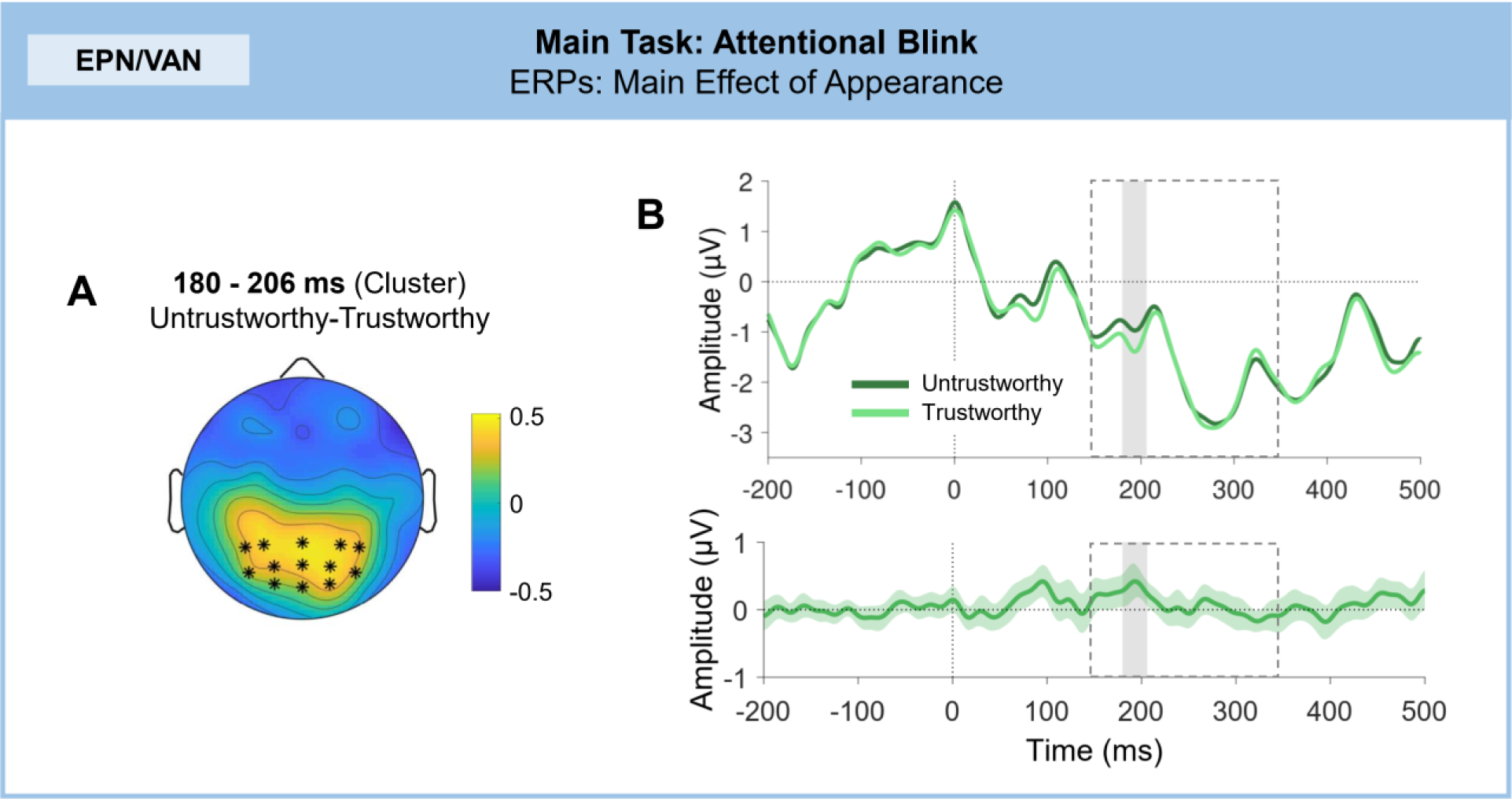
ERP differences in the attentional blink: Main effect of appearance. Cluster-based permutations tests revealed a significant difference between untrustworthy and trustworthy looking faces in the EPN time range, with enhanced positive amplitudes in the untrustworthy condition. (A) Difference topography for the appearance effect in the EPN cluster interval. Asterisks mark the electrodes included in the cluster. (B) Grand-average event-related potential for each appearance condition, based on pooled activity across electrode sites included in the cluster (upper panel), and corresponding difference curve with 95% bootstrap confidence interval (lower panel). The dashed frame indicates the time range included in the CBP test; the gray frame indicates the temporal extent of the cluster.

**Figure A5.2.**
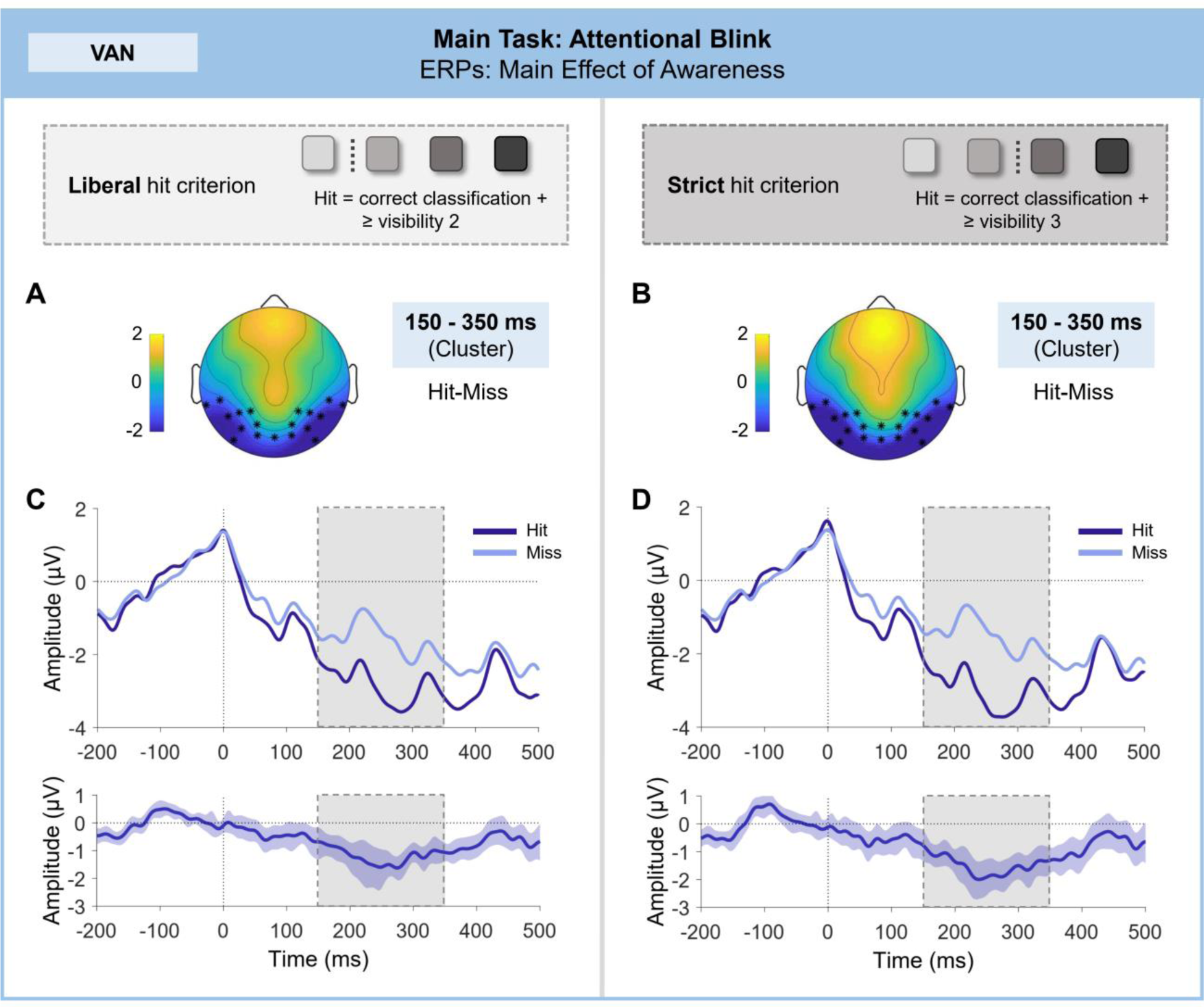
VAN differences in the attentional blink: Main effect of awareness. Cluster-based permutation tests revealed a significant main effect of awareness in the VAN, for both the liberal and strict hit criterion. (A), (B) Difference topography for the awareness effect in the VAN cluster interval. Asterisks mark the electrodes included in the cluster. (C), (D) Grand-average event-related potential for each awareness condition, based on pooled activity across electrode sites included in the cluster (upper panel), and corresponding difference curve with 95% bootstrap confidence interval (lower panel). The dashed frame indicates the time range included in the CBP test; the gray frame indicates the temporal extent of the cluster. As the observed cluster spanned the whole tested time range, both are identical here.

**Figure A5.3.**
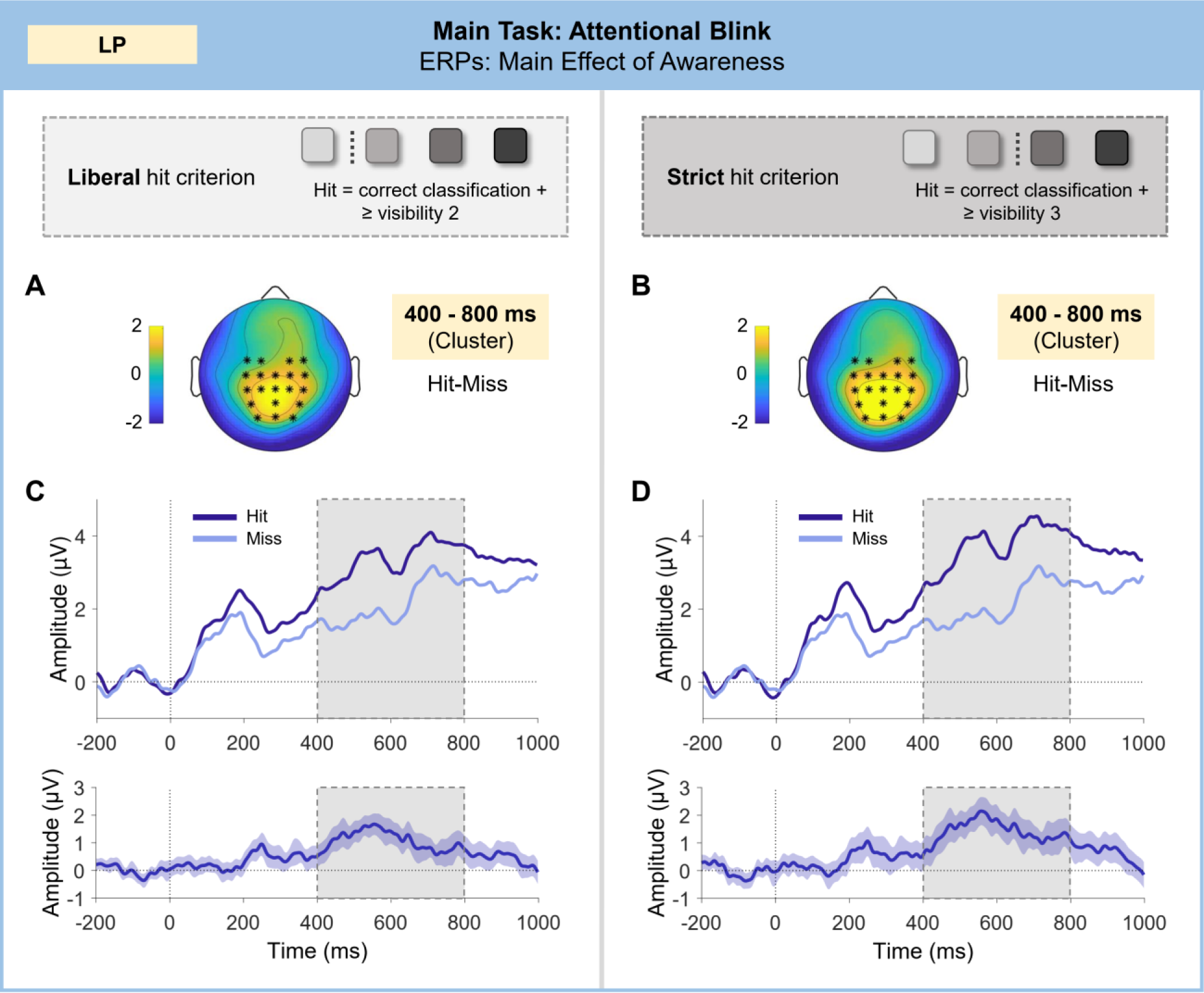
LP differences in the attentional blink: Main effect of awareness. Cluster-based permutation tests revealed a significant main effect of awareness in the LP, for both the liberal and strict hit criterion. (A), (B) Difference topography for the awareness effect in the LP cluster interval. Asterisks mark the electrodes included in the cluster. (C), (D) Grand-average event-related potential for each awareness condition, based on pooled activity across electrode sites included in the cluster (upper panel), and corresponding difference curve with 95% bootstrap confidence interval (lower panel). The dashed frame indicates the time range included in the CBP test; the gray frame indicates the temporal extent of the cluster. As the observed cluster spanned the whole tested time range, both are identical here.

